# Multispectral Imaging Differentiates Unique Macrophage Profiles in Patients with Distinct Chronic Liver Diseases

**DOI:** 10.1101/794610

**Authors:** Omar A. Saldarriaga, Adam L. Booth, Benjamin Freiberg, Jared Burks, Santhoshi Krishnan, Arvind Rao, Netanya Utay, Monique Ferguson, Minkyung Yi, Laura Beretta, Heather L. Stevenson

## Abstract

Intrahepatic macrophages influence the composition of the microenvironment, host immune response to liver injury, and development of fibrosis. Compared to stellate cells, the role of intrahepatic macrophages in the development of fibrosis remains ill defined. Multispectral imaging allows detection of multiple markers *in situ* in human formalin-fixed, paraffin-embedded tissue. This cutting-edge technology is ideal for analyzing human liver tissues since it allows spectral unmixing of fluorophore signals, subtraction of auto-fluorescence, and preserves architecture and the *in vivo* hepatic milieu. We analyzed resident Kupffer cells (CD68+), monocyte-derived macrophages (Mac387+), pro-fibrogenic macrophages (CD163+), and co-expression of pro-inflammatory (CD14) and anti-inflammatory (CD16) markers in liver biopsies from patients with hepatitis C virus (HCV) and different stages of fibrosis. Liver biopsies with advanced fibrosis showed increased accumulation of CD163+, MAC387+ and CD68+ macrophages in the portal tracts when compared to those with minimal fibrosis. Imaging software generated t-distributed stochastic neighbor embedding (t-SNE) plots and phenotype matrices that facilitated comparison of macrophage profiles. These included monocyte-derived (CD68+/Mac387+) and pro-fibrotic/anti-inflammatory (CD163+/CD16+) phenotypes. We established that the utility of this platform could be extended to liver biopsies from patients with other chronic liver diseases including nonalcoholic steatohepatitis and autoimmune hepatitis. Each disease exhibited a unique profile after spectral imaging analysis and this platform holds the potential to identify patients predisposed to progressive liver disease based on the macrophage composition. In summary, spectral imaging is a powerful tool that enables analysis of macrophage profiles in different types of chronic liver diseases and has potential to change the manner in which we evaluate liver biopsies leading to more personalized treatment strategies.

## INTRODUCTION

Under normal conditions, the liver maintains a tolerogenic microenvironment to avoid over activation of the immune response when it is exposed to a plethora of antigens that enter from the gut via the portal vein (1). One of the main cells thought to be a central player in the maintenance of hepatic tolerance is the resident hepatic macrophage, or embryonically derived Kupffer cell. Once this normally tolerogenic barrier is breeched, hepatic stellate cells differentiate into extra cellular matrix - producing myofibroblasts, initiating the path towards fibrosis development. Stellate cells have been extensively studied in the development of hepatic fibrosis and represent less than 1% of the non-parenchymal cells in the liver, while Kupffer cells make up greater than 20% (2). Once the tolerogenecity of the liver is lost and the hepatic microenvironment shifts to a more pro-inflammatory milieu, studies have shown that monocyte-derived macrophages are recruited to the liver (3–5). Primary human Kupffer cells exhibit a mixed expression of classical (M1) and alternative activated (M2) macrophage markers, with a dominant M2 phenotype that expresses CD163 (6, 7). CD68 and Mac387 identify human resident Kupffer cells and monocyte-derived macrophages, respectively (5, 8). Similar to flow cytometric studies of peripheral blood mononuclear cells (i.e., PBMCs), Liaskou et al. identified three critical monocyte-derived macrophage populations in human liver: pro-inflammatory or classical (CD14++/CD16-), anti-inflammatory or nonclassical (CD14+/CD16++) and the intermediate (CD14++/CD16+) type that perpetuate chronic inflammation and promote hepatic fibrosis (3, 9).

In the majority of patients, chronic HCV infection induces recruitment of immune cells into the liver, including monocytes/macrophages (6), and promotes an immunosuppressive environment that allows viral persistence (10, 11). HCV induces activation of signaling pathways that promote alternative macrophage activation (M2) and results in increased expression of pro- (M1) and anti-inflammatory (M2) cytokines (12, 13), which enhances overall hepatic inflammation and development of fibrosis by promoting differentiation of hepatic stellate cells (11, 13). However, it is now widely accepted that the “M1” and “M2” classification for macrophages is an over simplification since they exhibit extensive plasticity and change between homeostatic and pathologic conditions (14, 15).

Intrahepatic macrophages are of critical importance in the progression of hepatic inflammation and fibrosis in non-alcoholic steatohepatitis (NASH). NASH develops by multiple hits stemming from lipid accumulation, metabolic disruption, and oxidative stress in hepatocytes resulting in a pro-inflammatory state with subsequent activation of Kupffer cells and recruitment of monocyte-derived macrophages (16). Although less well understood, macrophages also play a role in development of autoimmune hepatitis (AIH) through their action as antigen presenting cells. Pro-inflammatory (M1) macrophages are increased in patients with AIH (17) and macrophage/Kupffer cell function is associated with increased fibrosis progression (18).

The inherent plasticity of macrophages renders *in vitro* systems suboptimal for study, since it is a challenge to faithfully replicate the hepatocyte microenvironment. Macrophages are difficult to isolate from human liver tissue and become activated, changing their phenotype when manipulated or cultured (19, 20). While flow cytometry is able to analyze multiple antigens on cellular suspensions of freshly isolated macrophages, it is unable to visualize them in the context of intact hepatic architecture (21) and fresh human tissue is not always available. Moreover, mouse models of certain liver diseases, such as those induced by HCV infection or NASH are poor surrogates since they fail to replicate the chronicity observed in humans (22). Routine immunohistochemical (IHC) staining of human liver biopsies can identify macrophages in formalin-fixed paraffin-embedded (FFPE) tissues; however, there are many limitations, including the inability to stain multiple antigens on the same cellular compartment (23), and depend on the availability of primary antibodies raised in different species to prevent cross-reactivity (21).

Recently, a cutting-edge technique has been developed that allows *in situ* characterization of human cells in FFPE tissues, the Vectra® 3 automated quantitative pathology imaging system. There are several reports using this technology, mainly for studying tumor infiltrating lymphocytes and cancer-related immunologic pathways (24, 25). This platform is ideal as it allows spectral unmixing of fluorophore signals with subtraction of background auto-fluorescence, producing a clean signal without interference from neighboring spectral wavelengths. Opal-conjugated fluorophores with tyramide signal amplification (TSA) are used to increase signal intensity and remove antibodies with each antigen retrieval (26). For this study, we hypothesized that this platform would successfully quantify and phenotype intrahepatic macrophages *in situ*. We optimized a 6-color multiplex immunofluorescence panel and spectral imaging analysis first in HCV+ patients with different stages of fibrosis and then applied this same approach to patients with different types of chronic liver disease (e.g., NASH and AIH).

## METHODS

### Patient liver biopsy specimens

The University of Texas Medical Branch Institutional Review Board approved the study protocol and all studies were conducted on de-identified, archived liver biopsies collected from 2006 to 2017. Liver biopsies were selected from healthy controls (n = 8) (i.e., liver biopsies from patients without known liver diseases and minimal histopathologic findings at the time of biopsy) and from patients with clinically (by serology and molecular testing) and biopsy confirmed HCV (with minimal fibrosis stage (n = 5) and advanced fibrosis (n = 6). The HCV+ and control patient groups were age, sex, BMI, and genotype matched. The majority of the patients were also HIV+. We also tested the platform in a patient with NASH (n = 1) and AIH (n = 1). Biopsies from all patients (Supplementary Table 1) were obtained as standard of care by licensed radiologists via the percutaneous route using an 18-gauge core needle. Tissue was immediately placed into 10% buffered formalin and processed using a TissueTek VIP tissue processor, paraffin-embedded, and sections were cut at 3 μm using Thermo-Fisher CryoStar NX70 cryostats. H&E and Masson’s trichrome stains were conducted on a Ventana Ultra automated stainer. All processing and staining was performed in a CAP-accredited histology laboratory by licensed histotechnologists.

### Fibrosis quantitation

Since all of the patient liver biopsies were collected as standard of care, the amount of inflammatory activity/injury and the stages of fibrosis had been previously determined by a board-certified pathologist (H.L.S) using a widely accepted scoring method (27). We used a quantitative method to determine the collagen proportionate area (i.e., CPA), similar to published reports (28). Briefly, each Masson’s trichrome stain was digitally scanned on an Aperio ImageScope Digital slide scanner and low power (1.2X) images of the entire surface area of the tissue core was measured in number of pixels using Nikon’s NIS Elements-BR Imaging Software. Next, the blue stained areas from the Masson’s trichrome stain were identified and specific thresholds were set using the software. The fibrotic area was divided by the total biopsy surface area to obtain the percent fibrosis (% fibrosis= (∑fibrotic area in pixels/∑ total area in pixels) x100). Liver capsule and large portal tracts (identified by the presence of nerve bundles) were excluded from these analyses.

### Antibody selection, monoplex assays, spectral library development and imaging analysis

The Vectra 3 platform (Akoya Biosciences, Hopkinton, MA), included inForm software, and Opal^TM^, 7-color manual IHC kit (Akoya Biosciences, Hopkinton, MA, OPAL^TM^ 7-Color Manual IHC kit, 50 slides, cat. Number: NEL811001KT) was used for optimization of the multiplex antibody panel. Unstained tissue sections from the above patient groups were used for these analyses and additional details are described in the supplementary methods (including suppl Table 2 and Figure 1).

**Figure 1.**
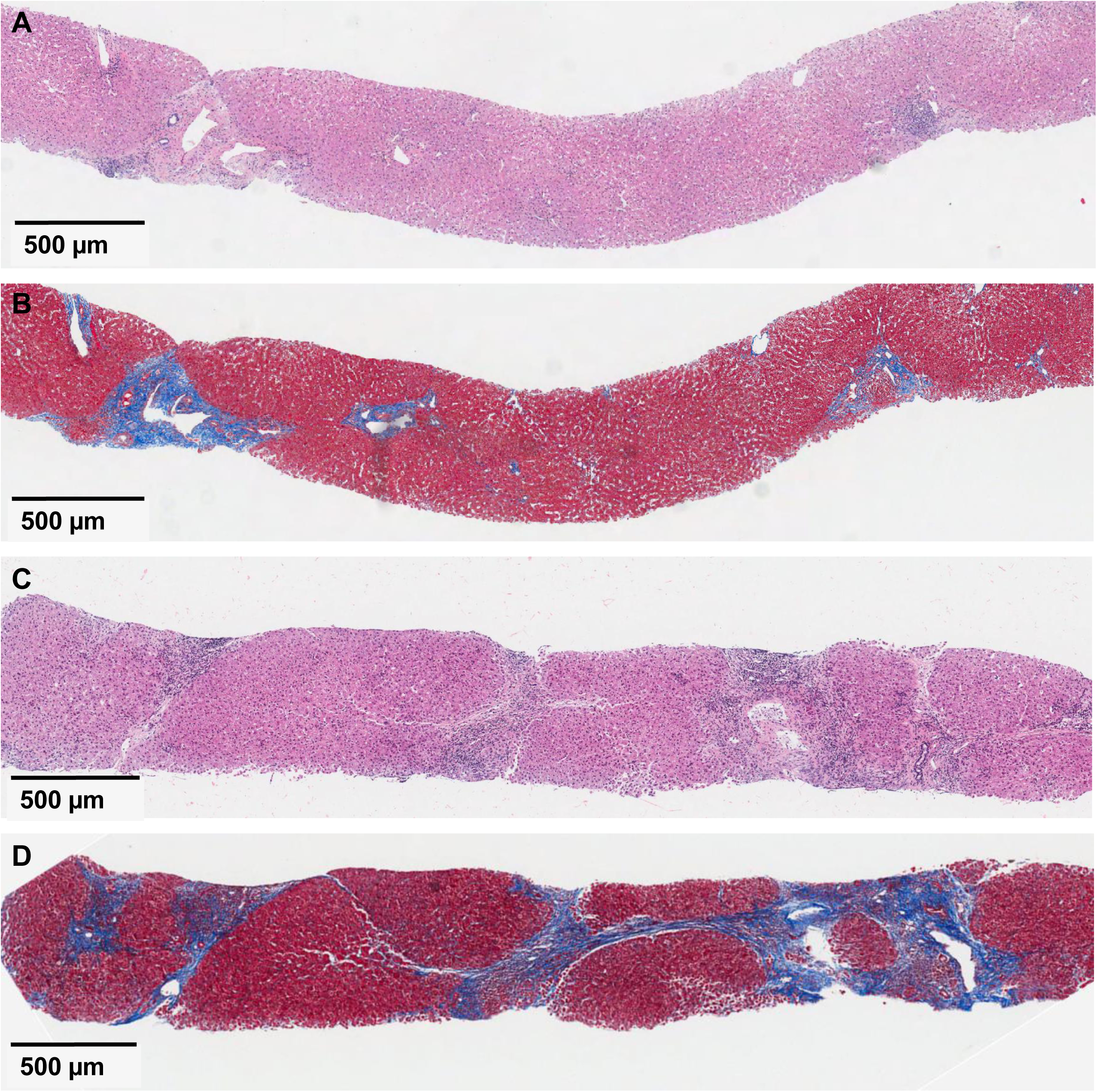
Histopathologic features of HCV with minimal and advanced fibrosis. Representative images from liver biopsies collected from patients with minimal and advanced stages of fibrosis. Compare images A (H&E) and B (Masson’s trichrome) that were obtained from an HCV+ patient with minimal fibrosis (Ishak stage: 1/6) to images C (H&E) and D (Masson’s trichrome) that were collected from an HCV+ patient with advanced fibrosis (Ishak fibrosis stage: 6/6).

## RESULTS

### Histopathologic features in HCV+ patients with minimal and advanced stages of fibrosis and quantification of collagen proportionate area

The initial optimization experiments analyzed liver biopsies from five HCV+ patients with minimal fibrosis (i.e., Ishak fibrosis stages: 1-2 out of 6) and six patients with advanced fibrosis (i.e., Ishak fibrosis stage: 5-6 out of 6). The two groups were age, gender, BMI, and genotype matched. Ishak criteria were used for scoring the liver biopsies as described in the materials and methods (27). Demographic information and clinical data were collected for all patients included in the study (Suppl Table 1). Figure 1 depicts the histopathologic features observed in patients with chronic HCV. A representative patient with minimal inflammatory activity and fibrosis (MHAI: 4/18; Fibrosis stage: 1/6) (Fig 1A and 1B) is compared to a representative patient with moderate inflammatory activity and advanced fibrosis (MHAI: 7/18; fibrosis stage: 6/6) (Fig 1C and 1D). The collagen proportionate area (CPA) was determined (28) and representative examples from liver biopsies with minimal (i.e., Ishak fibrosis stages: 1-2 out of 6) versus advanced fibrosis stages (i.e., Ishak fibrosis stage: 5-6 out of 6) are shown in Fig. 2. The overall median percentages of fibrosis were significantly different between the two groups (p < 0.01); patients with minimal fibrosis had a median of CPA of 5.299% (range: 1.613 to 8.021) and those with advanced liver fibrosis had a median CPA of 24.4% (range: 15.5 to 54.865)(**Suppl Table 1**).

**Figure 2.**
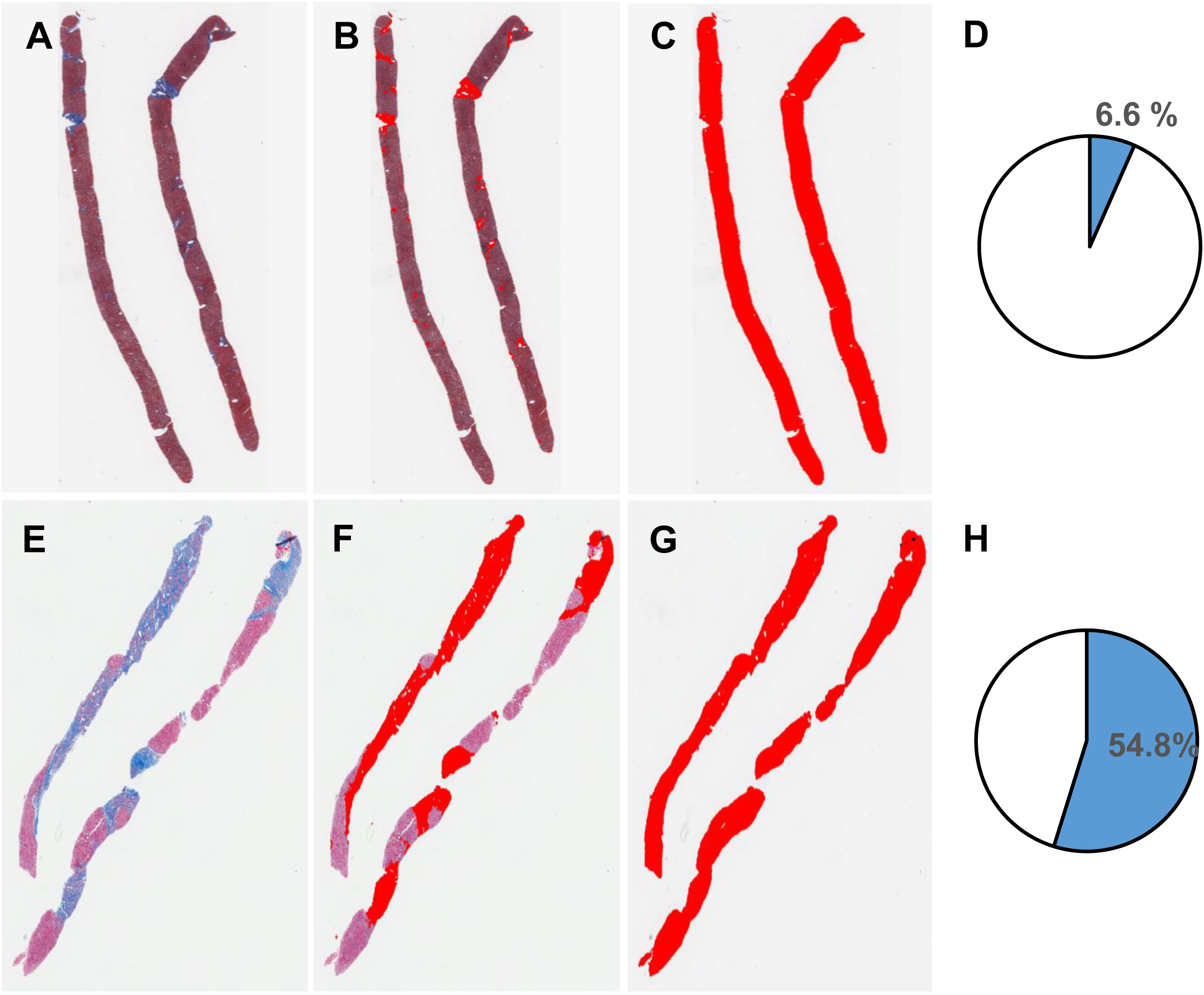
Determination of collagen proportionate area in HCV+ patients with minimal or advanced fibrosis using an advanced digital methodology. Slides from two representative HCV+ patients with either minimal (A-D; CPA: 6.6%) or advanced fibrosis (E-H; CPA: 54.8%) were stained with Masson’s trichrome (A,E; blue= fibrosis). Slides were scanned using an Aperio whole slide scanner and low power images (1.2X) were obtained using Aperio Image Scope software. Fibrotic regions or CPA (B,F) were determined by setting specific thresholds using NIS elements BR and divided by the total surface area of the biopsy (C,G) to calculate the percentage of fibrosis (D,H); (%CPA=∑fibrotic area/∑ total area).

### Monoplex assay optimization and spectral library development

An initial requirement for the standardization of this multiplex protocol was the evaluation of each macrophage marker singly in a “monoplex” assay in order to optimize the antibody/opal concentrations required to achieve the ideal morphologic staining pattern and proper target intensities (**Suppl Table 2**). AR6 buffer combined with use of the EZ retriever microwave system was effective in preventing non-specific staining. Individual monoplex stains without DAPI were used to create the spectral library, which provided the precise patterns for each fluorophore emission spectrum that was later required for spectral unmixing of multiplex stained slides (Suppl Fig 1). The staining pattern of the macrophage markers, which was evaluated by fluorescence (Suppl Fig 2, **Panel A**) and simulated bright field view using the inForm software (Suppl Fig 2, **Panel B**), were restricted to specific cellular compartments, as previously reported: CD68 (cytoplasmic), CD163 (cytoplasmic/membranous), CD14 (membranous), CD16 (membranous) and MAC387/S100A (nuclear/cytoplasmic) (3, 5). CD14 was also expressed on liver sinusoidal endothelial cells as previously reported (29, 30).

### HCV+ patients with advanced fibrosis had increased expansion and infiltration of intrahepatic macrophage populations

We evaluated the expression of five different macrophage markers including antibodies to identify resident Kupffer cells (CD68+), monocyte- or systemically-derived macrophages (Mac387+), anti-inflammatory or pro-fibrogenic (CD163+) macrophages, as well as common pro-inflammatory (CD14) and anti-inflammatory (CD16) markers. Their expression was evaluated in liver biopsies obtained from control patients (n = 8) and from HCV+ patients with minimal (n = 5) and advanced (n = 6) stages of fibrosis. Representative multiplex images from patients with minimal and advanced fibrosis, respectively, are shown for comparison (Fig 3A and 3B). The tissue segmentation feature of the inForm software was used to separate the liver biopsy tissue into portal tract and lobular regions (Fig. 3C and 3D). CD163+, MAC387+ and CD68+ macrophage expansion and infiltration (absolute number/area) was increased in the portal tracts of patients with advanced fibrosis when compared to those with minimal fibrosis (Fig. 3E). t-SNE analysis (31) uses dimensional reduction to allow visualization of cell similarities in two dimensional plots, and the multiplex panel was able to identify unique patterns of macrophages in the livers of patients without known liver disease (red), minimal fibrosis (blue) and advanced fibrosis (green) (Fig 3F).

**Figure 3.**
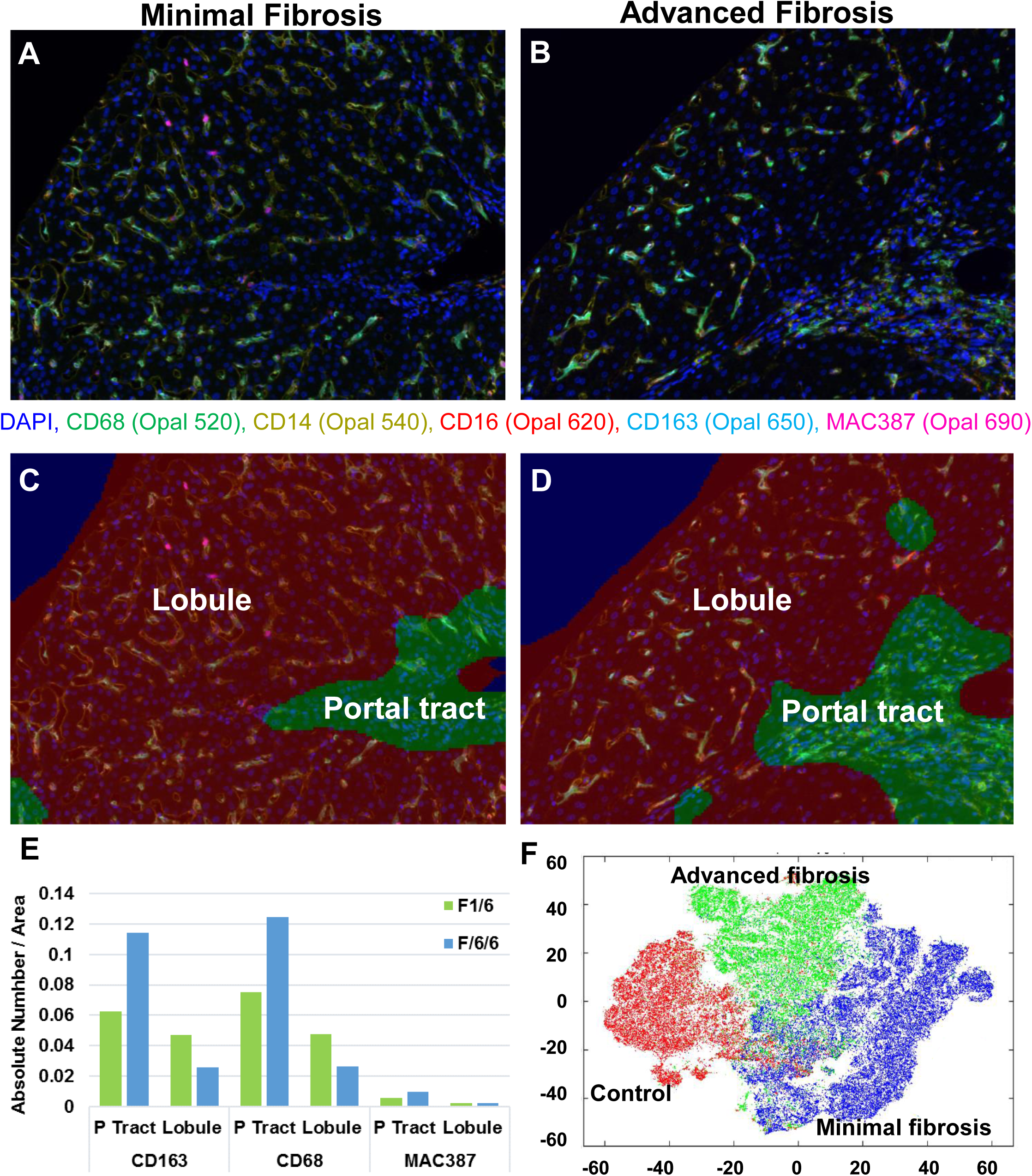
Intrahepatic macrophage populations are increased in HCV+ patients with advanced fibrosis when compared to those with minimal fibrosis. FFPE liver biopsies from HCV+ patients with minimal (n = 5) and advanced (n = 6) fibrosis were stained with the multiplex macrophage panel (see materials and methods). The Phenochart application was used to uniformly identify 50% of the biopsy tissue and multiple ROIs were acquired at 20X using the Vectra 3 platform. Representative images from these patients are shown in figures A-D. (A) Patients with minimal fibrosis showed decreased macrophage infiltration in the portal tracts and lobules, when compared to biopsies from patients with advanced fibrosis (B). (C) The analysis of images with Visiopharm applications revealed an increase in the absolute numbers of CD163+, MAC387+ and CD68+ macrophages in the portal tracts of patients with advanced fibrosis when compared to patients with minimal fibrosis. (D) t-SNE plots highlight the unique profiles of macrophages that were identified in the livers of patients with minimal (blue cluster) and advanced fibrosis (green cluster) compared to controls (red cluster).

### Unsupervised analysis of multiplex imaging data revealed significant differences in pro-inflammatory, anti-inflammatory, and monocyte-derived macrophage populations in patients with HCV-induced advanced hepatic fibrosis

We collected ROIs from at least 50% of each liver biopsy, which generated 30 to 50 multiplexed images per patient. Mean normalized data were further analyzed using a t-SNE plot of all the markers involved from all the cells of all sample groups (Fig 4A) and on a per-group basis (Fig 4B). The proportion of total macrophages calculated in controls was 20.9%, minimal fibrosis was 17.3%, and advanced fibrosis was 22.6%. We identified five unique macrophage clusters and overlaid them onto their corresponding t-SNE plots (Fig 4C), which showed significantly different expression patterns between controls and patients with advanced fibrosis. Specifically, the proportions of CD68+CD14+CD163+ macrophages and the overall expression of CD14+ clusters were significantly increased in control livers when compared to patients with advanced fibrosis (Fig 4D, **#1 and #2, respectively**). Although the proportion of CD14+ cells in the patients with minimal fibrosis was not significantly different when compared the controls or patients with advanced fibrosis, it was also the population with the highest frequency in this group. Liver biopsies from patients with advanced fibrosis revealed three distinct macrophage populations with significantly increased cell numbers: 1) CD68-CD14-CD163+CD16+Mac387-, 2) CD68+CD14-CD163-CD16-Mac387-, and 3) CD68+CD14-CD163-CD16-Mac387+ (Fig 4D, **#3, #4 and #5, respectively**).

**Figure 4.**
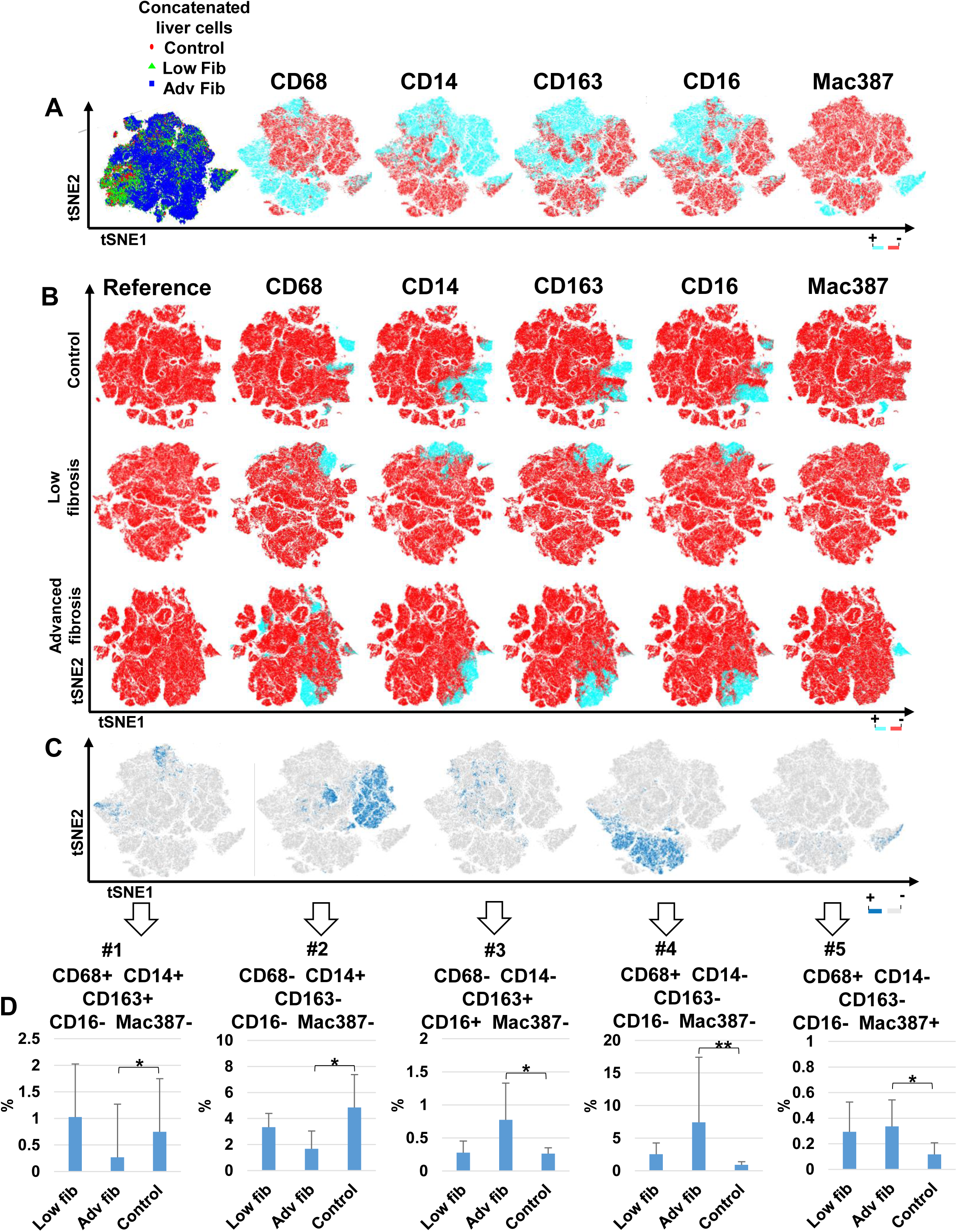
Expression heat maps from t-SNE and phenotype cluster analyses of control and HCV+ livers with minimal and advanced fibrosis. Multispectral imaging data of hepatic macrophage from human liver biopsies: controls (n = 8), minimal fibrosis (n = 5) and advanced fibrosis (n = 6), were analyzed with a dimensional reduction t-SNE algorithm. Heat maps showing the positive (opal) or negative (red) expression of all macrophage markers (CD68, CD14, CD163, CD16 and Mac387) in the concatenated t-SNE (A) and individually within the control, minimal fibrosis and advanced fibrosis groups (B). Five PhenoGraph clusters that were significantly increased between control and HCV advanced fibrosis were overlaid individually on the concatenated t-SNE plot from the figure 4A. Clusters of cells expressing the different markers, positive (blue) or negative (gray) are shown (C). Frequency of five macrophage subpopulations significantly increased in control and advanced fibrotic livers (D). All data presented as median, with significance calculated using the Wilcoxon rank sum test (*p < 0.05; ***p < 0.01).

### Liver biopsies from human patients with different types of chronic liver disease have distinct macrophage phenotypes

We then assessed this platform’s capacity to delineate other chronic liver diseases. We used the same macrophage multiplex panel followed by imaging analysis to compare liver biopsies from control patients versus those collected from patients with HCV, NASH and AIH. Fig. 5 compares representative multiplex images acquired from each group (**Panel A**) and examples of tissue segmentation analyses (**Panel B**). Each disease displayed a unique pattern of portal and lobular macrophage populations based on the morphology observed in the multiplex images (Fig 5, **Panel A**). When compared to controls, liver biopsies collected from patients with chronic liver diseases showed not only increased numbers of macrophages in the portal tracts, but reflective of more activated-appearing macrophages within the lobules (Fig 5, **Panels A and B**). All patients with chronic liver disease showed increased Mac387+ monocyte-derived macrophages in the portal tracts when compared to controls; however, the numbers were markedly increased in NASH and AIH (Fig 5, **Panel C**). A liver biopsy from one patient with severely active AIH (MHAI: 13/18; Ishak fibrosis stage: 2/6) revealed striking increases in both portal and lobular macrophages when compared to controls (Figure 5). Analysis of each channel individually in this patient with AIH (Suppl. Fig 2) confirmed the existence of unique populations for each parent macrophage marker (i.e., CD68, CD163, and Mac387). Although the distribution of the CD68+ and CD163+ macrophages looked similar, the CD163+ staining pattern was noticeably more intense consistent with the presence of larger macrophages. Multiple foci within the lobules identified areas devoid of obvious hepatocytes (i.e., “hepatocyte drop-out”) that instead comprised large aggregates of macrophages (Suppl Fig 2, **Panels A and B;** Fig 5, **Panel A, AIH**). These areas correlated with the aggregates of pigment/ceroid-laden macrophages detected by light microscopy in patients with active NASH or AIH (H&E and PASD stains, Fig 5 and Suppl Fig 2, **insets**).

**Figure 5.**
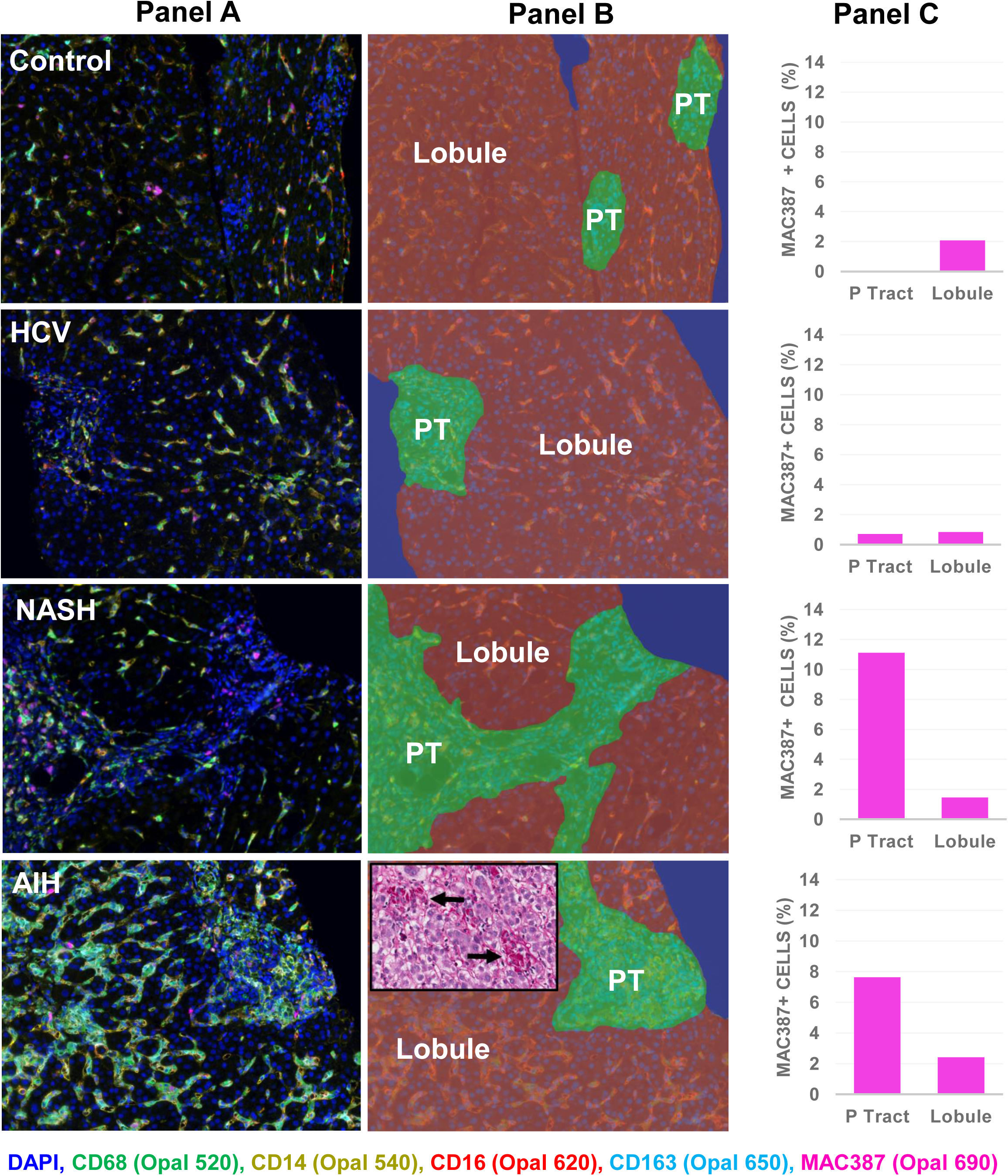
Spectral imaging analysis of patients with different types of chronic liver diseases (HCV, NASH and AIH) highlights the complexity of intrahepatic macrophage phenotypes in a single 20X multiplex image.. **Panel A**: Fluorescent multispectral composite unmixed images of different liver diseases stained with the macrophage panel (see materials and methods). **Panel B**: Manual tissue segmentation using InForm; portal tract (green), lobule (red), other (blue). (20X images). **Panel C**: Shows differences in the frequency of Mac387+ macrophages in the portal tracts and lobules in patients with either HCV, NASH or AIH, compared to control patients. Patients with NASH and AIH have increased MAC387+ macrophages within their portal tracts when compared to patients with HCV or controls. The aggregates of ceroid/pigment-laden macrophages that are often observed with liver injury (an example from the patient with AIH is shown in the inset, arrows), including that induced by AIH and NASH, expressed CD163 with high intensity (also see **Suppl Fig 2**). Results shown are representative of three different patients with similar hepatitis activity scores (i.e., MHAI) and fibrosis stages (using the Ishak criteria) (48). AIH: Autoimmune hepatitis; HCV: hepatitis C virus, PT: Portal tract.

The distinct macrophage phenotypes present in the hepatic microenvironment and the variation between different types of chronic liver disease became apparent when visualizing the cell phenotyping data (Fig 6, **Panel A**), where each color represents a specific macrophage subtype. By creating unique phenotypic maps and t-SNE plots for the various types of chronic liver diseases, we could more easily visualize the unique macrophage profiles in these patients. In the t-SNE plots, healthy controls showed approximately six different tightly clustered macrophage populations with less cellular diversity when compared to patients with chronic liver diseases. The latter exhibited marked increases in the breadth of cellular phenotypes and expansion of different populations (i.e., the populations were less tightly clustered) (Fig 6, **Panel B**).

**Figure 6.**
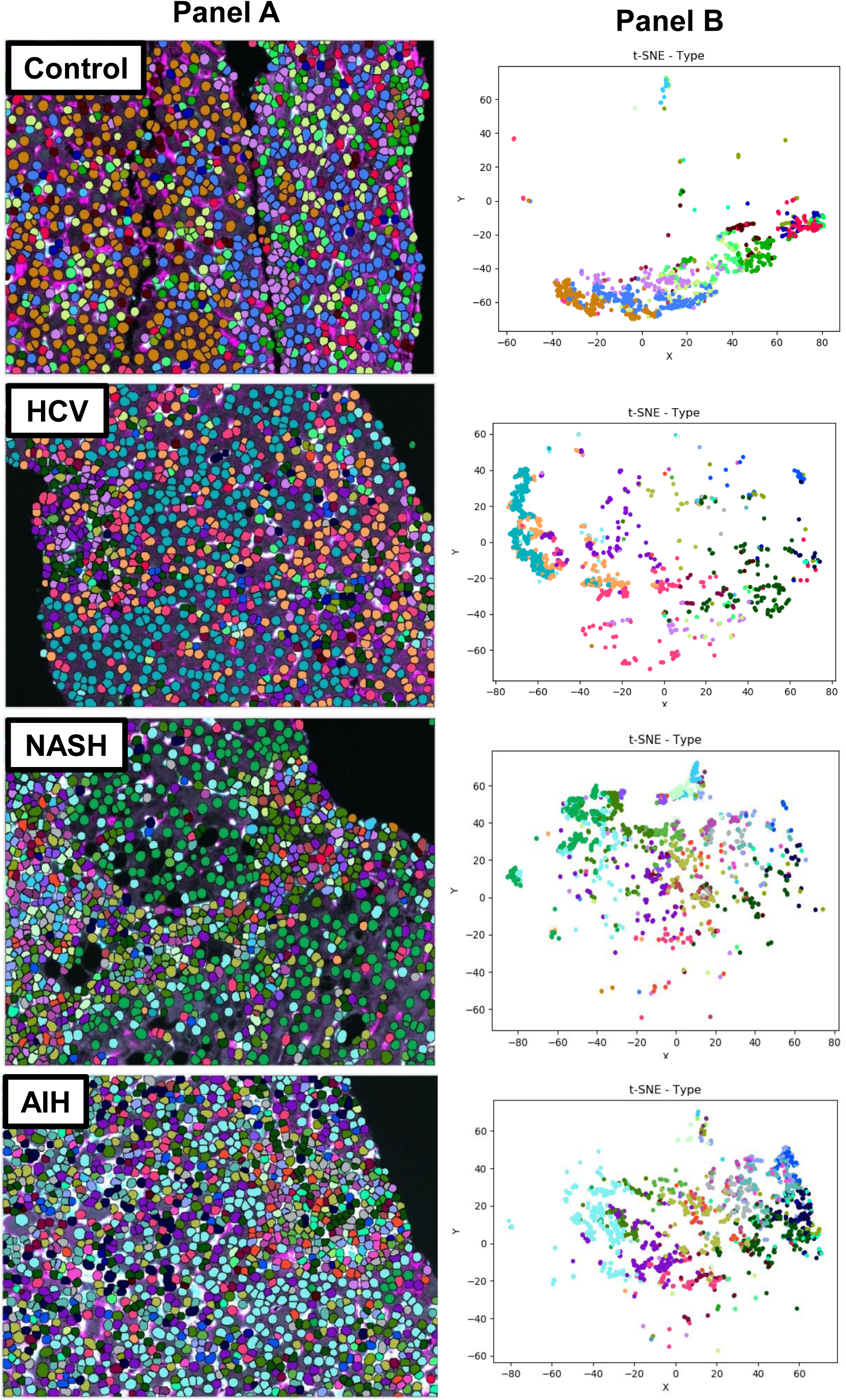
Cell phenotyping images and t-SNE plots highlight diverse macrophage populations in a single multiplex liver biopsy image acquired from patients with different chronic liver diseases (HCV, NASH and AIH). The slides were stained with the macrophage multiplex panel and representative single multiplex images (20X) were selected and acquired. The multiplex images were then analyzed with Visiopharm phenotyping applications and each color represents a unique cellular phenotype (**Panel A**). The t-SNE plots (**Panel B**) highlight the unique patterns of concatenated macrophage markers that are present in the livers of patient with chronic liver diseases versus controls. Cells with similar properties appear close together in the two-dimensional map and red (or “hot”) markers show cells with relatively more expression of that specific marker when compared to blue (or “cold”) markers, which indicate absent or minimal expression. Control patients showed much less diversity in the types of macrophages and other cellular phenotypes when compared to patients with chronic liver disease. Results shown are representative of three different patients with similar hepatitis activity scores and fibrosis stages. AIH: Autoimmune hepatitis; HCV: hepatitis C virus; NASH: non-alcoholic steatohepatitis.

We next wanted to determine the number of unique macrophage phenotypes present in each of the multiplex images (shown in Fig 5, **Panel A and** Fig 6, **Panel A**) from patients with chronic liver diseases versus controls, using Visiopharm phenotype matrix algorithms. For both the phenotypic matrices and t-SNE plots (Fig 7 and Suppl. Fig 3-6), red (or “hot”) markers denote cells with enhanced expression of the indicated marker when compared to blue (or “cold”) markers, representing minimal or undetectable expression. The phenotype matrix algorithm identified unique macrophage profiles in different types of chronic liver disease and images from these patients showed increased complexity and more numerous cellular phenotypes when compared to controls. To clarify these observations, the phenotype matrix plot generated from the multiplex image representing a control patient’s liver biopsy identified approximately nine different cellular phenotypes corresponding to a distinct macrophage lineage (all CD68+, CD163+, or Mac387+ phenotypes: Fig 7, **#8-10, 12, 14-17, 18).** The phenotypes that are all blue (i.e., negative for expression of all markers, e.g., #1-7), most likely represent hepatocytes or other cell types that fail to express any of the macrophage markers in the multiplex panel. We did not include the phenotype that only showed expression of CD14+ (Fig 7, **#13**); since this antigen may not be sufficiently macrophage-specific and identify other cell types including sinusoidal endothelial cells. It is noteworthy that patients with chronic liver diseases exhibited an increased diversity of macrophage phenotypes with approximately 19, 17, and 12 different types observed in HCV+, NASH, and AIH, respectively (Fig 7, amongst the phenotypes positive for CD68 and/or CD163 and/or Mac387). The corresponding t-SNE plots derived from the images are shown in the supplementary figures and again highlight the increased cellular complexity observed in patients with chronic liver disease when compared to controls (Suppl Fig 3-6).

**Figure 7.**
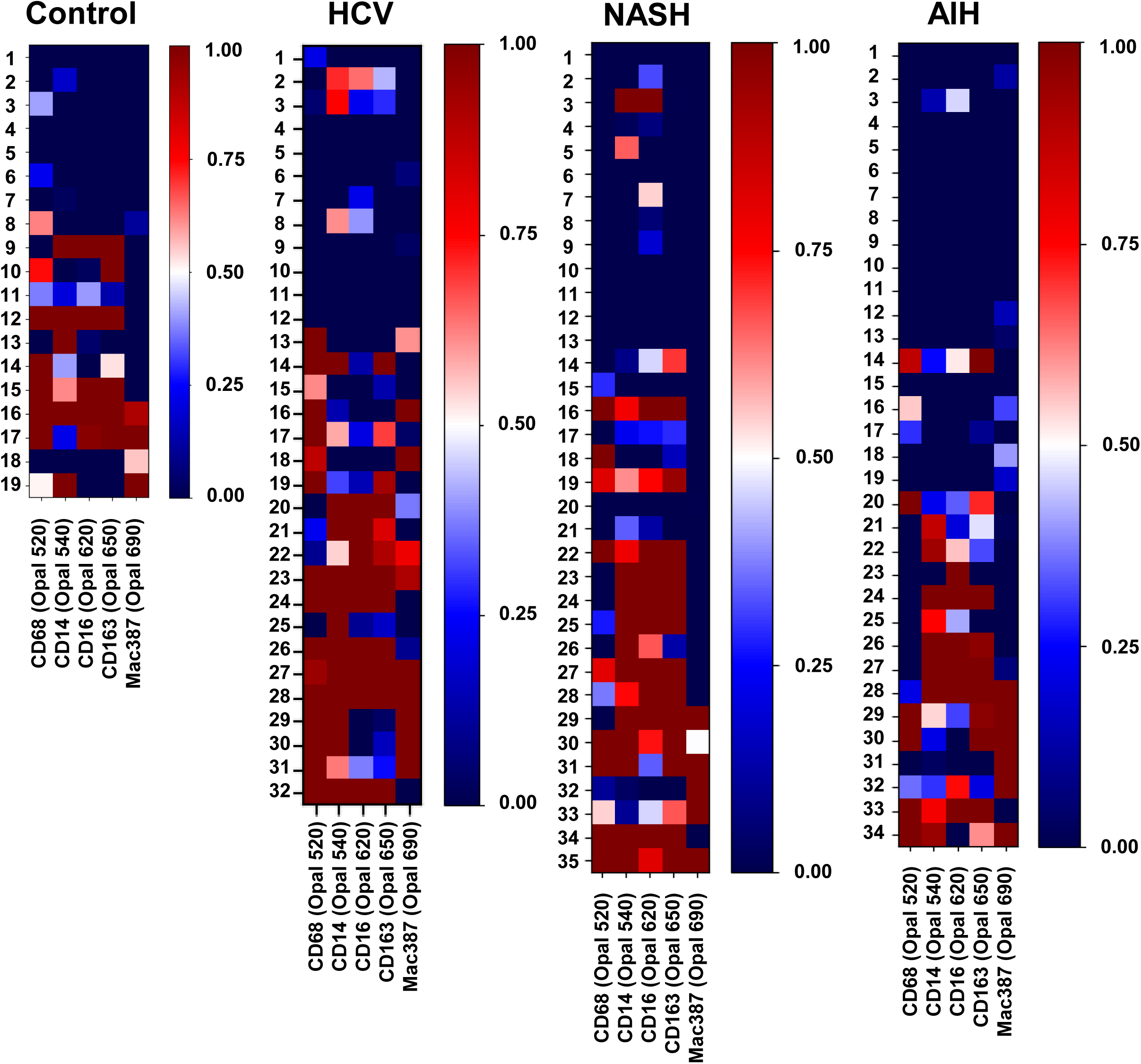
The phenotypic matrix algorithm identified expansion in the numbers and unique macrophage phenotypes in liver biopsies obtained from patients with chronic liver disease when compared to controls. Visiopharm phenotype matrix algorithms were used to determine the different phenotypes identified in the single 20X images acquired from liver biopsies collected from controls and patients with either HCV, NASH, or AIH as shown in Figures 5 and 6. Comparison of the phenotypes that expressed either CD68, CD163 or Mac387, showed that the number and complexity of the different phenotypes increased in the diseased liver biopsies when compared to controls (Controls: at least 9; HCV: at least 19; NASH: at least 17; AIH: at least 12). The CD14+/CD16+ phenotypes also likely represents a macrophage population. Results shown are representative of three different patients with similar hepatitis activity scores and fibrosis stages. AIH: Autoimmune hepatitis; HCV: hepatitis C virus; NASH: non-alcoholic steatohepatitis.

Phenotype matrix analyses exposed the presence of several different phenotypes that co-expressed CD68+ and Mac387+ in HCV+ patients (Fig 4D, Fig 7 **phenotypes #16, 18, 23, 27-31**). Monocyte-derived cells that were predominantly positive for Mac387 only, were present in patients with NASH (Fig 7 **phenotypes #32, and #33**) and AIH (Fig 7, **phenotypes #31, 32**). CD68+ and CD163+ macrophage populations were not identical in each diseased state and certain cells expressed only one of these markers. Unique populations of CD163+CD68- or CD163-CD68+ macrophages were identified (Fig 7: **controls, phenotypes #8 and #9; HCV, phenotypes #15, 16, 20-22, 29-31; NASH, phenotypes #18 and #23-25; and AIH, phenotypes #26-28)**. t-SNE plots were also generated for individual markers in controls and in each type of chronic liver disease (Suppl Fig. 3-6).

## DISCUSSION

Studying macrophages *in situ* in human liver biopsy tissue, where they naturally reside, offers a more meaningful approach that will provide an in depth understanding of their role in the complex hepatic microenvironment. Macrophages present in the portal tracts and lobules are not the same (32) and have unique phenotypes and densities that varied between controls, patients with minimal fibrosis, patients with advanced fibrosis, and different chronic liver diseases, as shown in this study. There are many advantages to using a spectral imaging platform: 1) multiple antigens can be detected simultaneously, even on the same cell or within the same cellular compartment; 2) it eliminates impediment of spectral overlap and the auto-fluorescence inherently present in liver tissue; 3) it can be conducted on FFPE tissues, a common biospecimen source; and 4) it allows *in situ* characterization of cells within the context of liver architecture and the preserved hepatic microenvironment. These attributes are of even greater importance when studying cells such as macrophages that are notorious for extreme plasticity (33, 34) and become activated when they are manipulated (19, 20). While *in vitro* methods have provided us with much insight regarding the biology of these mercurial cells, perturbations concomitant with *in vitro* procedures likely served to mask phenotypes and functions present in the human liver.

A critical step for successful multiplex staining with this panel is proper preservation of FFPE biopsy material. We recommend using freshly cut unstained slides stored at room temperature if used within one week, or alternatively stored at -80°C and protected from moisture for longer periods. For the initial monoplex stains, optimal staining was determined empirically by evaluating different antibody concentrations followed by adjusting the concentration of Opal-TSA to obtain target intensities.

Likewise, the staining sequence for the antibodies in the multiplex reaction and the use of different AR buffers (i.e., AR6 or AR9) proved to be critical steps in optimizing the multiplex panel (**suppl Table 2**).

Since infiltrating macrophages are known to be increased in patients with HCV (5, 35), we wanted to validate our multiplex macrophage panel imaging modality using this disease. We compared differences in the numbers of CD163+, CD68+, and Mac387+ cells in liver biopsies collected from patients with HCV and observed increases in all of these phenotypes in the portal tracts of patients with advanced fibrosis (Fig 3E) similar to previous reports (32). Kupffer cells comprise ~20% of the non-parenchymal cells in the liver (36), and since the number of Mac387+ cells was minimal when compared to the numbers of CD68+ cells, we conclude that the majority of CD68+ cells detected are Kupffer cells. Increased numbers of resident CD68+ macrophages in both portal tracts and lobules and increased numbers of circulation-derived macrophages (e.g., identified by Mac387+/ S100A9+ in this study) within the portal tracts have been reported in patients with HCV (5) and was confirmed with the spectral imaging platform. Recruitment into the liver of circulation-derived macrophages results in increased pro-inflammatory cytokine expression including IL1-β and IL6, culminating in a persistent pro-inflammatory state, activation of resident Kupffer cells and subsequently hepatic stellate cells (5).

The 6-color panel used for these analyses was able to show distinct differences between the cell populations identified in controls, patients with minimal fibrosis, and patients with advanced fibrosis (Fig 3F). Imaging analysis using the customized algorithm revealed significant increases in two cell clusters in liver biopsies from patients without known liver disease, specifically a population of CD68+/CD14+ macrophages and importantly, overall increased expression of CD14 (Fig 4D, **#1,#2**). The identification of a CD68+CD14+ cell population thought to have pro-inflammatory, immunoregulatory, and patrolling type functions, similar to previous findings in tolerogenic livers, and a recent study of human intrahepatic macrophages that analyzed these populations with single-cell RNA sequencing (7, 15). This study also confirmed the accumulation of different macrophage populations (Suppl Fig 4) that were previously shown to be increased in patients with chronic HCV or in human monocytes cultured with HCV-infected hepatocytes (5, 12, 13, 32, 37). Three main macrophage clusters that were significantly increased in the patients with advanced fibrosis included CD68+ macrophages, CD68+Mac387+ monocyte-derived macrophages, and CD163+CD16+ anti-inflammatory macrophages (Fig 4D, **#3,#4,#5).** The results of these imaging analyses confirm previous observations that suggest a predominance of CD14+ cells during homeostasis and increased accumulation of CD16+CD163+ macrophages in human liver diseases (3, 6, 11, 13). The overall decrease in CD14 in patients with advanced fibrosis when compared to controls is likely due to lower expression of CD14 not only on macrophages, but also on liver sinusoidal endothelial cells, which are also known to express this antigen and have been shown to decrease expression upon activation, such as that that occurs with HCV infection (29).

Interpretation of CD163+ macrophage function has varied depending on the study; some reports support a tolerogenic phenotype with high basal expression in non-diseased liver, while other studies suggest they confer a characteristic “M2” phenotype that is increased in portal tracts and lobules of patients with chronic HCV (11, 13, 32). This study reveals a more nuanced result where several macrophage subtypes expressing this antigen are present in liver biopsies from control patients; however, the number of CD163+ macrophage phenotypes increased concomitant with disease state (Fig 7). The intermediate pro-inflammatory monocyte subset CD14++/ CD16+, which accumulates in the chronically inflamed human liver, also expresses CD163 and this phenotype is thought to represent a pro-fibrogenic macrophage (3, 9). Many of the patients included in this study were co-infected with HIV (73%) (**Suppl Table 1**) and these patients were determined to have significantly higher expression of CD163 on CD14++CD16+ monocytes when compared to healthy individuals (38). We recently showed that HIV infection of macrophages did not influence the fibrogenic gene expression of hepatic stellate cells, but potentiated the HCV-mediated fibrogenic activation of hepatic stellate cells (39).

The images obtained using Visiopharm immune phenotyping applications following staining with the multiplex macrophage panel, showed unique profiles indicative of the chronic liver disease type (Fig 5 and 6) and highlighted the complexity of the diseased hepatic microenvironment *in vivo* (Fig 5-7 and Suppl Fig 3-5). The phenotype matrix algorithm revealed unique phenotypes, where certain macrophage populations appeared to express high levels of CD68 with minimal CD163 (compare the red or “hot” populations of cells in the Opal 520 channel that are not present in Opal 650; Fig 7). In contrast, other populations expressed high levels of CD163 with minimal CD68 (Fig 7), which suggests that CD68+ and CD163+ macrophage populations are varied with diverse expression patterns by these markers. Amongst the three chronic liver diseases compared, biopsies obtained from patients with NASH were typified by highest infiltration of Mac387+ monocyte-derived macrophages in the portal tracts, and to a lesser extent AIH (Fig 5). However, given that some patient biopsies representing the minimal fibrosis group had increased numbers of Mac387+ cells in their portal tracts, prompts us to speculate that these samples came from patients predisposed to increased fibrosis. These data complement previous reports, by showing that chronic liver disease is associated with an increased prevalence of all types of macrophages including classical (CD14++/CD16-), intermediate (CD14++/CD16+) and non-classical (CD14+ or dim/CD16++), which are functionally thought to be pro-inflammatory, pro-fibrotic, and anti-inflammatory, respectively (3, 22). Liaskou et al. showed that the intermediate type preferentially accumulates in the inflamed liver and has phagocytic ability, secretes inflammatory cytokines (e.g., IL-1, IL-6, and TNF-alpha) and profibrogenic chemokines (e.g. CCL2 and CCL5), and purportedly plays a role in fibrosis development. We detected aggregates of CD163+/CD14++/CD16+ macrophages within the sinusoids in patients with AIH or NASH (Suppl Fig 2 and Fig 5) in close proximity to areas of subsinusoidal fibrosis (data not shown). The phenotyped multiplex images and t-SNE plots acquired in liver sections representing different types of chronic liver diseases highlighted the diversity of the populations detectable *in situ* under non-tolerogenic states (Fig 6).

The involvement of hepatic macrophages in disease severity and fibrosis development is most widely accepted in NASH, where it is known that Kupffer cells become activated to a more pro-inflammatory phenotype that results in increased inflammation, production of pro-inflammatory cytokines, recruitment of monocyte-derived macrophages into both the portal tracts and lobules, and activation of Stellate cells into myofibroblasts (40, 41). We showed that in contrast to controls, biopsies from patients with NASH or AIH showed increased Mac387+ monocyte-derived macrophages in portal tracts and developed large aggregates of CD163+ macrophages in the lobules (Fig 5 and Suppl. Fig 2). The CD163+ population correlated with the presence of frequent lobular ceroid-laden macrophages in the PAS-D stain by light microscopy (Fig 5 and Suppl Fig 2).

This study, limited to five different markers, established that categorizing macrophages into “M1” and “M2” phenotypes is much too simplistic. As we showed, a single image acquired from a liver biopsy from patients with HCV, AIH or NASH, revealed disease-specific profiles (qualitative) with up to 19 unique macrophage phenotypes (Fig 7), each depicted by a range of marker expression levels (quantitative) identified within the multiplex panel (Fig 6). We are one of the first to demonstrate the efficacy of this platform in studies of macrophages in non-neoplastic liver tissue, including in patients with HCV, NASH, and AIH.

Patients undergo an invasive procedure to obtain liver biopsy material. Instead of merely providing a diagnosis, as clinicians and pathologists we should glean as much information as possible from this precious material. In the future, imaging analysis platforms may inform precision medicine by personalizing treatments for chronic liver diseases such as NASH, AIH, and HCV based on the macrophage profile in the liver. We already have several treatments that target macrophages that are FDA approved (42) and the dual CCR2/CCR5 inhibitor that decreases monocyte recruitment to the liver is now in phase 3 clinical trials (40, 43). Keep in mind, even though the CENTAUR trial showed improvement in fibrosis in patients treated with this drug, only 20% responded (versus 10% in the placebo group); we need to determine how the macrophages in the remaining 80% of patients contribute to disease progression. Results of multiplex imaging platforms may allow us to personalize treatment in patients depending on the phenotypes expressed in their livers, similar to the approaches that determine phenotypic expression on tumor infiltrating lymphocytes in patients with cancer (24, 25, 44–47). The biggest challenge is for us to develop more rapid and efficient big-data analytics that can cope with the plethora of data generated. In summary, use of multispectral imaging has the potential to not only change our understanding of intrahepatic macrophages, but also the way in which patient liver biopsies are evaluated in the future.

## List of Abbreviations

AIH: autoimmune hepatitis
FFPE: formalin-fixed, paraffin-embedded
H&E: Hematoxylin and eosin
HCV: hepatitis C virus
IHC: immunohistochemical
LSEC: liver sinusoidal endothelial cells
MHAI: Modified hepatitis activity index
NASH: nonalcoholic steatohepatitis
PASD: Periodic acid–Schiff with diastase
ROI: regions of interest
TSA: tyramide signal amplification
t-SNE: t-distributed stochastic neighbor embedding

## ACKNOWLEDGEMENTS

We would like to thank Ben Freiberg from Visiopharm for his assistance and expertise in using the software. Bradley Dye and Judy Pham assisted with fibrosis quantification of liver biopsies. Thank you also to Kevin Mottershead from Akoya Biosciences for his support and guidance in using the Vectra Platform. We extend sincere gratitude to Jeffrey East, P.A., and Shana White, study coordinator, for their assistance with acquiring and organizing the clinical samples. Sam Diaz de Leon and Blanca Flores provided secretarial support and assistance with obtaining the archived liver biopsy tissue blocks.

## SUPPLEMENTARY METHODS

### Antibody selection, monoplex assays, and spectral library development

Tissue blocks were stored at room temperature and sections were cut immediately prior to multiplex staining, whenever possible. Slides that were unable to be stained within in one week were stored at - 80°C in slide storage boxes wrapped with Parafilm (to reduce exposure to moisture), which is recommended for antigen preservation in FFPE tissue sections (29). The Vectra 3 platform (Akoya Biosciences, Hopkinton, MA), included inForm software, and Opal^TM^, 7-color manual IHC kit (Akoya Biosciences, Hopkinton, MA, OPAL^TM^ 7-Color Manual IHC kit, 50 slides, cat. Number: NEL811001KT) was used for optimization of the multiplex antibody panel. Controls, including “monoplex” stains with each individual antibody/Opal fluorophore combination (without DAPI), DAPI alone, and an unstained liver biopsy tissue section, were used to build the spectral library for subsequent use during spectral unmixing in the multiplex assay (31). We performed titration of the primary antibody concentrations: CD68 (hepatic macrophage or Kupffer cell marker), Mac387 (systemic monocyte-derived macrophage marker), CD163 (established anti-inflammatoery or pro-fibrogenic macrophage marker), CD14 (inflammatory) and CD16 (anti-inflammatory) (**Suppl Table 2**). The fluorescence and simulated IHC views with inForm 3.1 software and synthetic library spectra were used to evaluate for proper cell morphology and staining pattern (evaluated by a subspecialty trained, board certified pathologist: H.L.S.). In order for the auto-fluorescence spectrum inherently present in liver tissue to be subtracted from future analyses, an additional unstained control slide was incubated with primary antibody, without the addition of Opal TSA and DAPI. Once the optimal staining condition for each antibody was obtained, Opal TSA concentrations were adjusted to achieve target intensities (5-30 inform normalized counts). We then selected the best antigen retrieval buffer (AR6, Biogenex, Fremont, CA versus AR9, Akoya Biosciences (Supplementary Table 2) to eliminate non-specific staining. Representative fields from the single color slides were imaged at 20x using the Vectra 3 and the spectra were extracted from the acquired images using inForm 3.1 and saved to the Spectral Library. The quality of the Spectral Library was assessed by evaluating unmixed images to confirm the absence of spectral overlap or bleed over between channels, and by evaluating the values of the unmixed peaks for each fluorophore (Suppl Fig 1).

### Opal multiplex assay development

Multiplex staining requires utilization of the optimized conditions determined from the monoplex slides, as described in the above section. The detection of five macrophage markers in human liver FFPE tissue samples was performed using the Opal^TM^, 7-color manual IHC kit. Slides were heated at 60°C for 45 min to 1 hour; then residual paraffin was removed using xylene (Fisher, Fair Lawn, NJ) (3x 10min), and tissue was rehydrated in a graded series of histological grade ethanol solutions (100% - 1x 10 min; 95% - 1 x 10 min; and rinse in 70%) and rinsed with distilled water. Slides were fixed in 10% neutral buffered formalin (Fisher, Kalamazoo, MI) for 30-45min, rinsed in water and placed in the standardized antigen retrieval buffers (ie., AR9 PE for CD68; **Suppl Table 2**). Slides were then microwaved at 95°C for 15 min using the EZ retriever system V3-110V (Biogenex, Fremont, CA), which uses a temperature controlled antigen retrieval system to solve the inconsistencies of different antigen retrieval methods. The slides were allowed to cool to room temperature and were then washed with water and tris-buffered saline Tween20 (TBST-0.1%). A hydrophobic barrier pen (Vector labs, Burlingame, CA) was used to surround the tissue section on the slide. Blocking was performed with 2-3 drops of antibody diluent/blocking solution (included with the Opal kit) for 10 min at room temperature in a humidified chamber, followed by incubation with 150µL of the first primary antibody (**Suppl Table 2**). After primary antibody incubation, the slides were rinsed in TBST and washed in TBST (3 x 2 min). Slides were incubated with 2 drops of the secondary antibody mix (polymer HRP Ms + Rb) for 10min. The slides were incubated for 10 min with 150µL of the standardized dilution of the assigned OPAL selected for the antibody in the position 1 (i.e., opal 520 (1:300) for CD68). After each antigen retrieval step the protocol restarts at the blocking step until all antibodies are used. AR6 buffer was always used before nuclear counterstaining with DAPI. Nuclei were stained with 150µL of DAPI (Akoya Biosciences) diluted in TBST (15µL/ mL X 4 min). After nuclear staining, the slides were washed for 2 min in TBST and 2 min in water before mounting with Invitrogen™ ProLong™ Diamond Antifade Mountant (Thermo Fisher Scientific, Grand Island, NY) and cover slipped. Samples were stored at 4°C until imaged. Whole stained slides were acquired using the Vectra 3 platform and phenochart software (Akoya Biosciences®). Fifty percent of the total biopsy tissue was uniformly identified for regions of interest (ROI) and captured using the 20X lens. Additional regions of portal tracts were captured to adequately capture a statistically relevant sampling of portal tracts due to their small size. Once we determined the absence of interference between each antibody signal and rebalanced specific signals to ensure that target intensities were in range, we proceeded to imaging analysis.

### Imaging analysis

For the HCV+ and control patient biopsies, we collected regions of interest (ROIs) from at least 50% of the tissue. Multiplexed images were unmixed, segmented and scored using inForm software. Thresholds for specific markers were adjusted using images from control patients and then applied to all the images using a batch analysis approach, processing only two markers at a time. Mean normalized counts from the different macrophage markers were overlapped per group. The heterogeneity and distribution of the various macrophage phenotypes in the study groups were determined using t-SNE and phenotypic matrix algorithms, which facilitated visualization and comparison of macrophage profiles in different liver biopsies, fibrosis stages, and disease types. The non-parametric Wilcoxon rank sum test (also called Wilcoxon–Man-Whitney test) was performed to compare all the possible marker combinations between the three patient groups (control, minimal and advanced fibrosis). Component TIFF files were exported and analyzed using Visiopharm® software. Different algorithm-based applications were developed using this platform, which identified specific marker thresholds to calculate the absolute numbers of parental macrophage populations (CD68, CD163 and MAC387) and the co-expression of specific markers (CD14, CD16). The tissue segmentation feature of the software was used to separate the hepatic parenchyma into portal tracts and lobules so that the number of cells could be enumerated in these regions. The heterogeneity and distribution of the various macrophage phenotypes in one single image from biopsies of patients with different liver diseases (HCV, NASH and AIH), was also determined using t-SNE and phenotypic matrix algorithms to confirm that the macrophage multiplex panel and Vectra platform could be applied easily to other types of chronic liver disease. Cell phenotype population differences between groups was compared using Wilcoxon Rank-Sum test with p <0.05 chosen for significance. The associated calculations and plotting were conducted using MATLAB 2019a (MATLAB 2019; *version 9.6.0, R2019a*, Natick, Massachusetts: The MathWorks Inc.).

**Supplementary Table 1.** Clinical data of HCV patients with minimal (ID: 1-5) and advanced fibrosis (ID: 6-11) included in this study. Abbreviations: ALT (alanine transaminase); AST (aspartate transaminase), HIV (Human immunodeficiency virus), HBV (hepatitis B virus), CPA (collagen proportionate area). *Pretreatment laboratory values.

| Patient | Age | Sex | ALT*<br>(U/L) | AST*<br>(U/L) | ALP*<br>(U/L) | Bilirubin*<br>(uMol/L) | Albumin*<br>(g/L) | HCVgenotype/<br>HIV / HBV | Ishak<br>Fibrosis<br>Stage | Fibrosis<br>(CPA) % |
| --- | --- | --- | --- | --- | --- | --- | --- | --- | --- | --- |
| 1 | 45 | F | 352 | 361 | 140 | 0.8 | 4.1 | +1a/+/- | 2 | 5.299 |
| 2 | 48 | M | 82 | 70 | 128 | 0.3 | 4.2 | +1a/+/- | 1 | 1.613 |
| 3 | 38 | M | 40 | 29 | 108 | 0.5 | 4.4 | +1a/+/- | 1 | 6.661 |
| 4 | 61 | M | 28 | 30 | 123 | 0.7 | 4.1 | +1a/+/- | 1 | 3.677 |
| 5 | 57 | M | 15 | 23 | 99 | 0.5 | 4 | +2b/+/- | 2 | 8.021 |
| 6 | 47 | F | 122 | 133 | 187 | 0.8 | 4.3 | +1b/+/+ | 5 | 15.664 |
| 7 | 63 | M | 91 | 93 | 72 | 1.4 | 3.8 | +1a/-/- | 5 | 24.419 |
| 8 | 53 | M | 48 | 85 | 165 | 2.0 | 3.3 | +1a/-/- | 6 | 54.865 |
| 9 | 43 | M | 111 | 89 | 99 | 4.8 | 0.4 | +1a/+/- | 5 | 31.681 |
| 10 | 48 | M | 51 | 46 | 120 | 0.7 | 4.7 | +1a/+/- | 5 | 15.5 |
| 11 | 63 | F | 239 | 168 | 94 | 1 | 4.3 | +2/-/- | 6 | 26.74 |

**Supplementary Table 2.** Antibodies and optimized multiplex conditions used to identify human macrophage populations in human FFPE liver biopsies. *Akoya Biosciences; **Biogenex, ***Ready to Use.

| **Antibody** | **Vendor/Clone/Isotype** | **Dilution** | **Incubation**<br>**Time**<br>**(min)** | **Antigen**<br>**Retrieval**<br>**Buffer** | **Opal**<br>**/Dilution** | **Position**<br>**in**<br>**Reaction** |
| --- | --- | --- | --- | --- | --- | --- |
| **CD68** | Biogenex/KP1/mouse-IgG1κ | RTU*** | 60 | AR9 (PE*) | 520/1:300 | 1 |
| **MAC387** | Dako/Mac387/mouse-IgG1κ | 1:200 | 30 | AR6 (Bio**) | 690/1:100 | 2 |
| **CD16** | Abcam/EPR16784/rabbit-IgG | 1:500 | 60 | AR6 (Bio**) | 620/1:600 | 3 |
| **CD14** | Abcam/EPR3653/rabbit-IgG | 1:500 | 45 | AR6 (Bio**) | 540/1:600 | 4 |
| **CD163** | Leica/10D6/mouse-IgG1 | RTU*** | 30 | AR6 (Bio**) | 650/1:100 | 5 |
| **DAPI** | Perkin Elmer | 15uL/mL | 4 | AR6 (Bio**) | - | 6 |

**Supplementary Figure 1.**
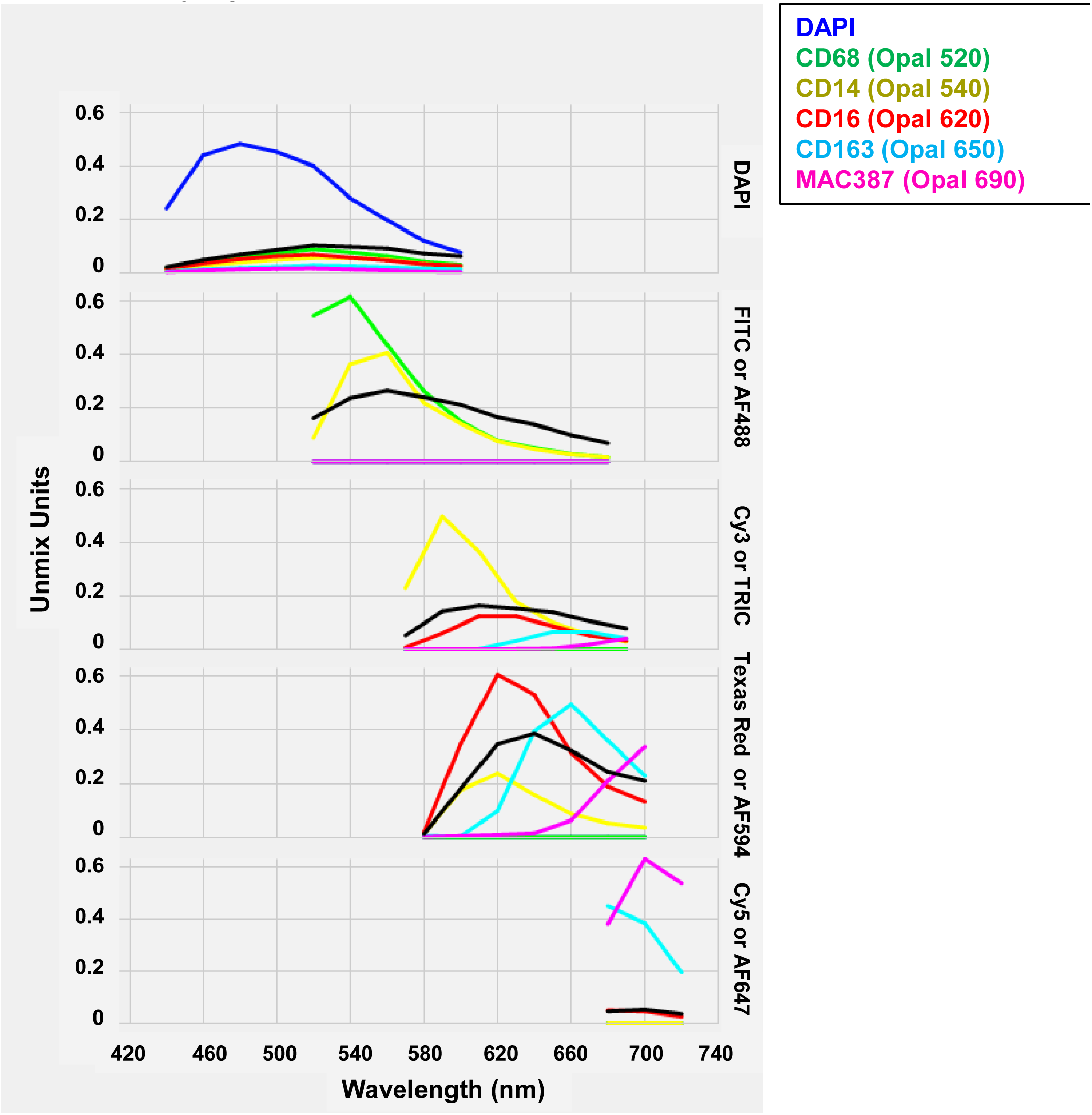
Spectral library generated from optimized “monoplex” stains and an unstained liver biopsy slide. A single slide from each antibody-OPAL fluorophore pair, a single slide with DAPI alone, and an unstained slide, were analyzed using inForm software (included with Vectra 3 platform) to create a “Spectral library.” The Spectral library is one of the main advantages of using the Vectra 3 platform as it allows elimination of spectral overlap by “unmixing” the various fluorophore signals and allows subtraction of autofluorescence.

**Supplementary Figure 2.**
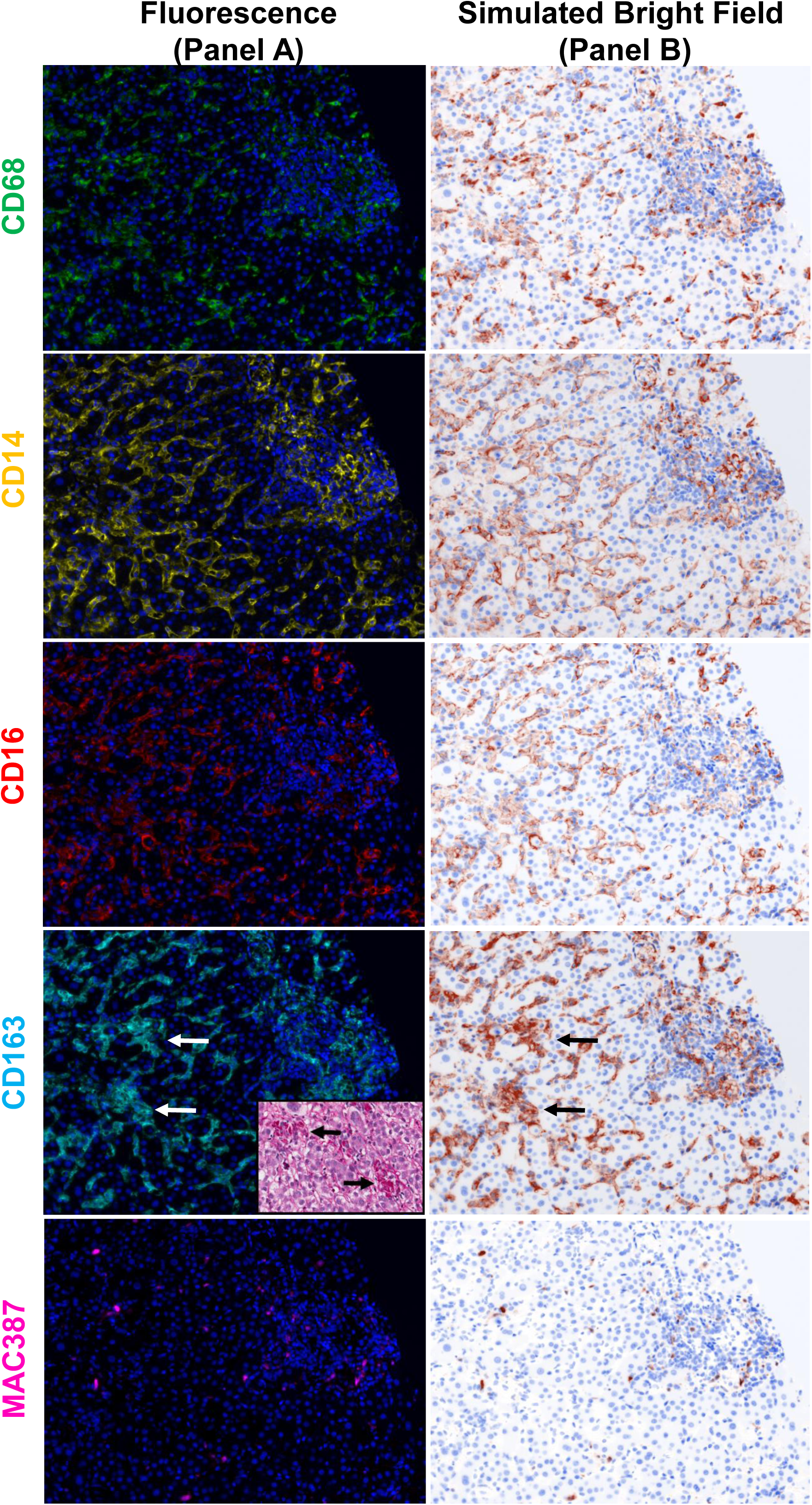
Visualization of each fluorophore channel acquired from multiplex staining a liver biopsy from a patient with AIH. **Panel A**: A representative fluorescence image obtained from the multiplex macrophage panel after spectral unmixing was used to separately view the expression of each individual macrophage marker. CD68 (green-Opal 520), CD14 (yellow-Opal 540), CD16 (red-Opal 620), CD163 (cyan-Opal650), MAC387 (magenta-Opal 690) and nuclear stain (Blue-DAPI). **Panel B**: Simulated brightfield images were also generated to recreate the staining pattern that would be observed by conventional IHC chromogenic methods. (20X images). Large aggregates of ceroid-laden macrophages were observed in the lobules and these showed high expression of CD163, which correlated with large aggregates or pigment/ceroid-laden macrophages by histology (inset; PASD stain).

**Supplementary Figure 3-6.**
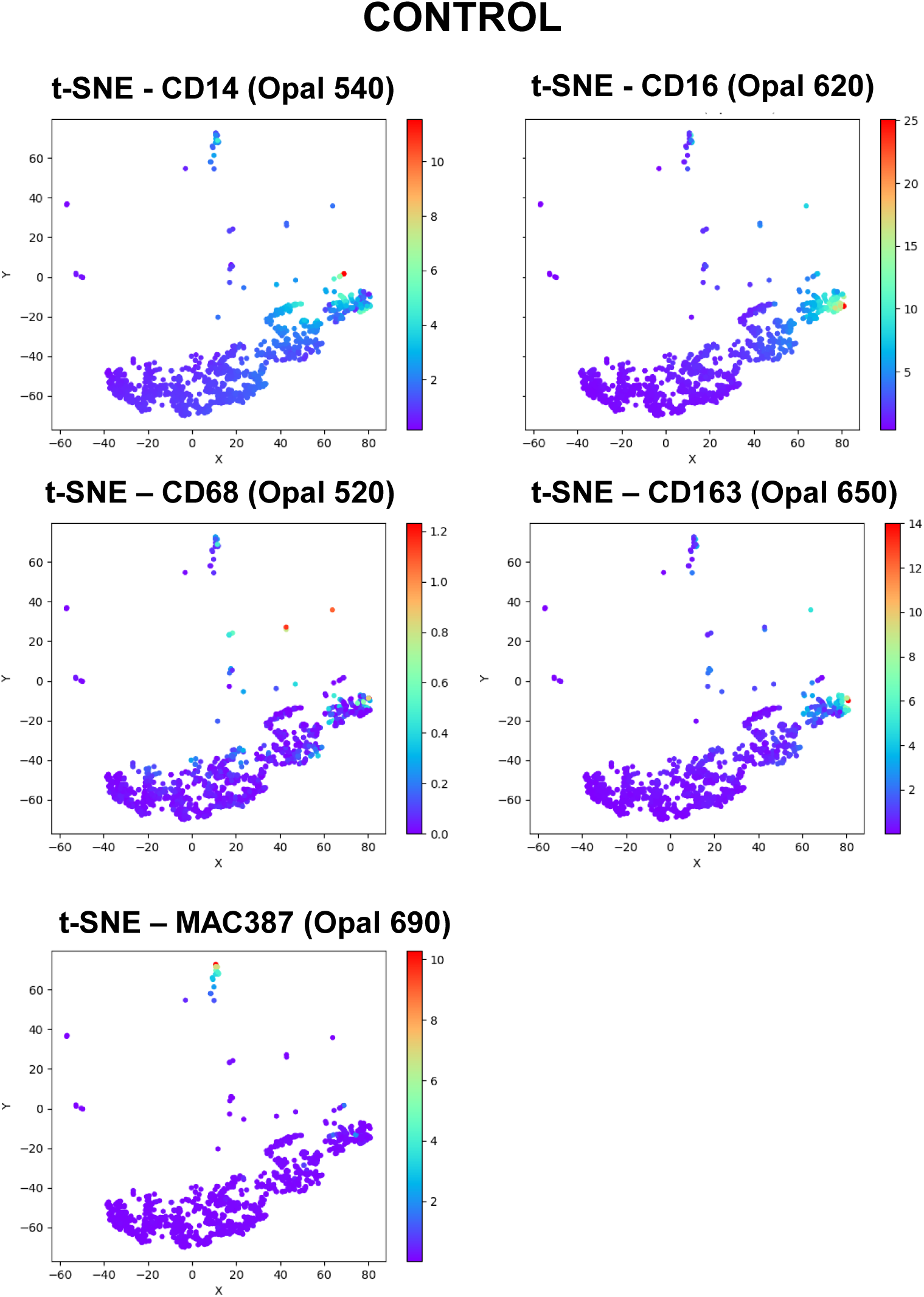
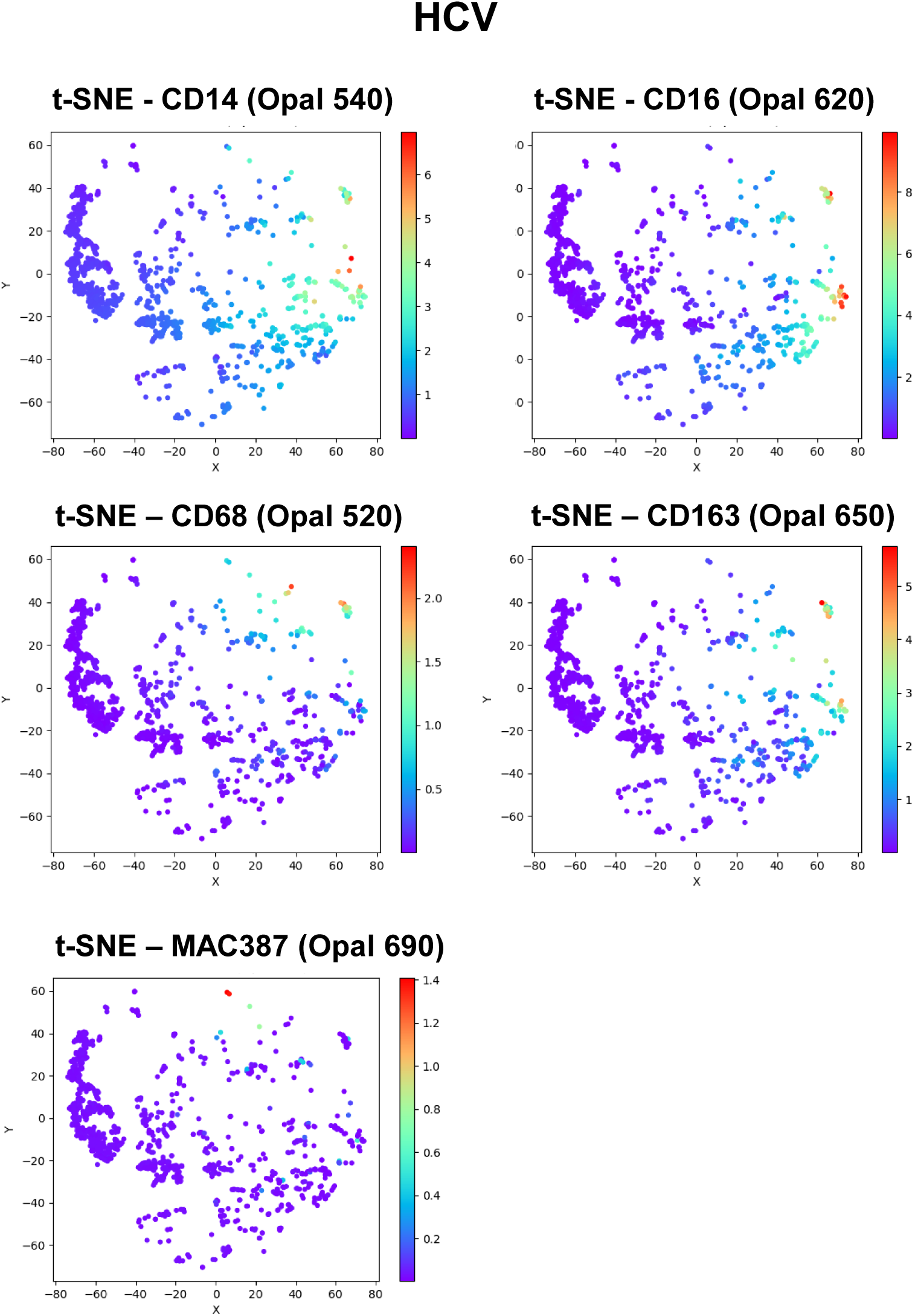
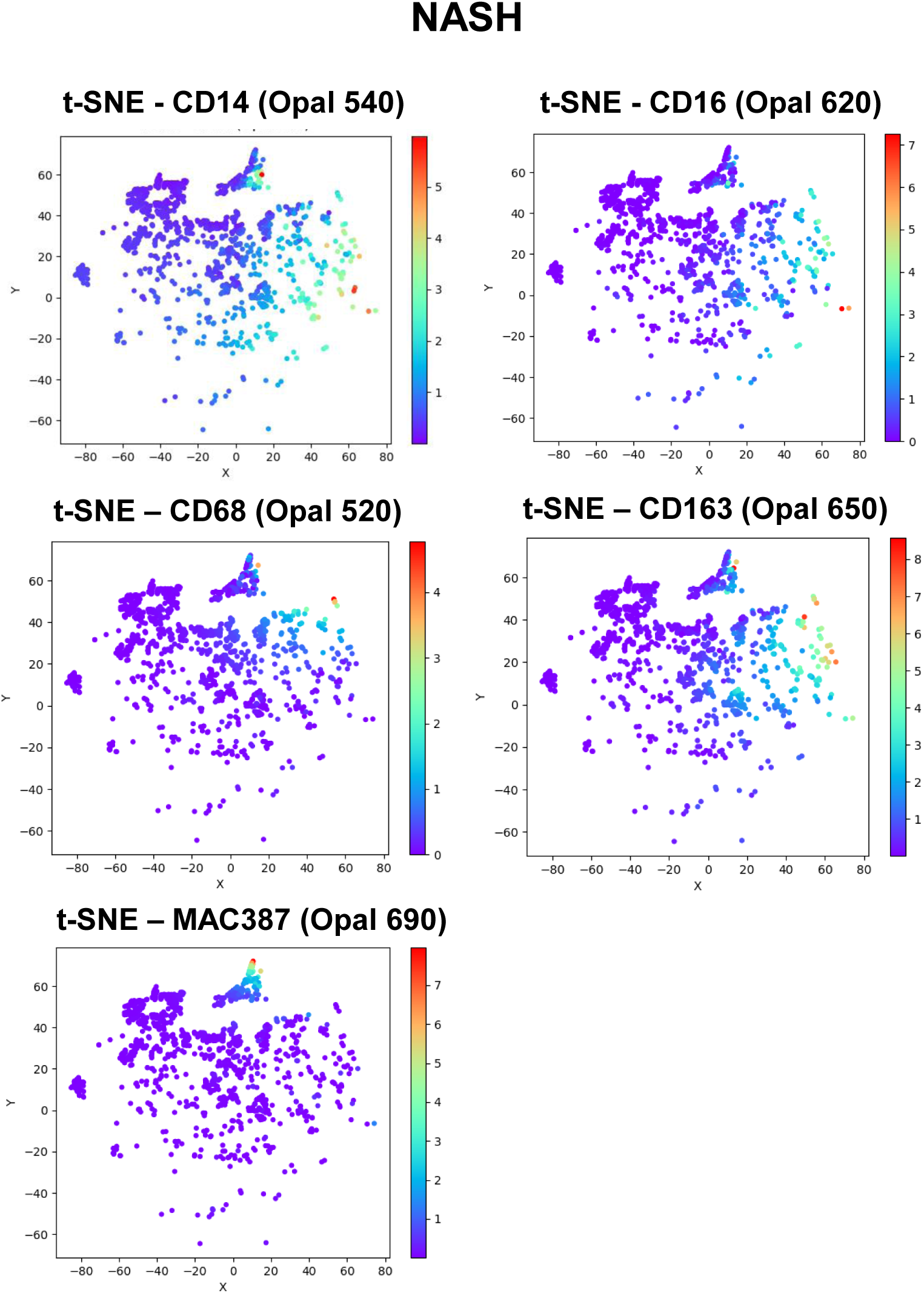
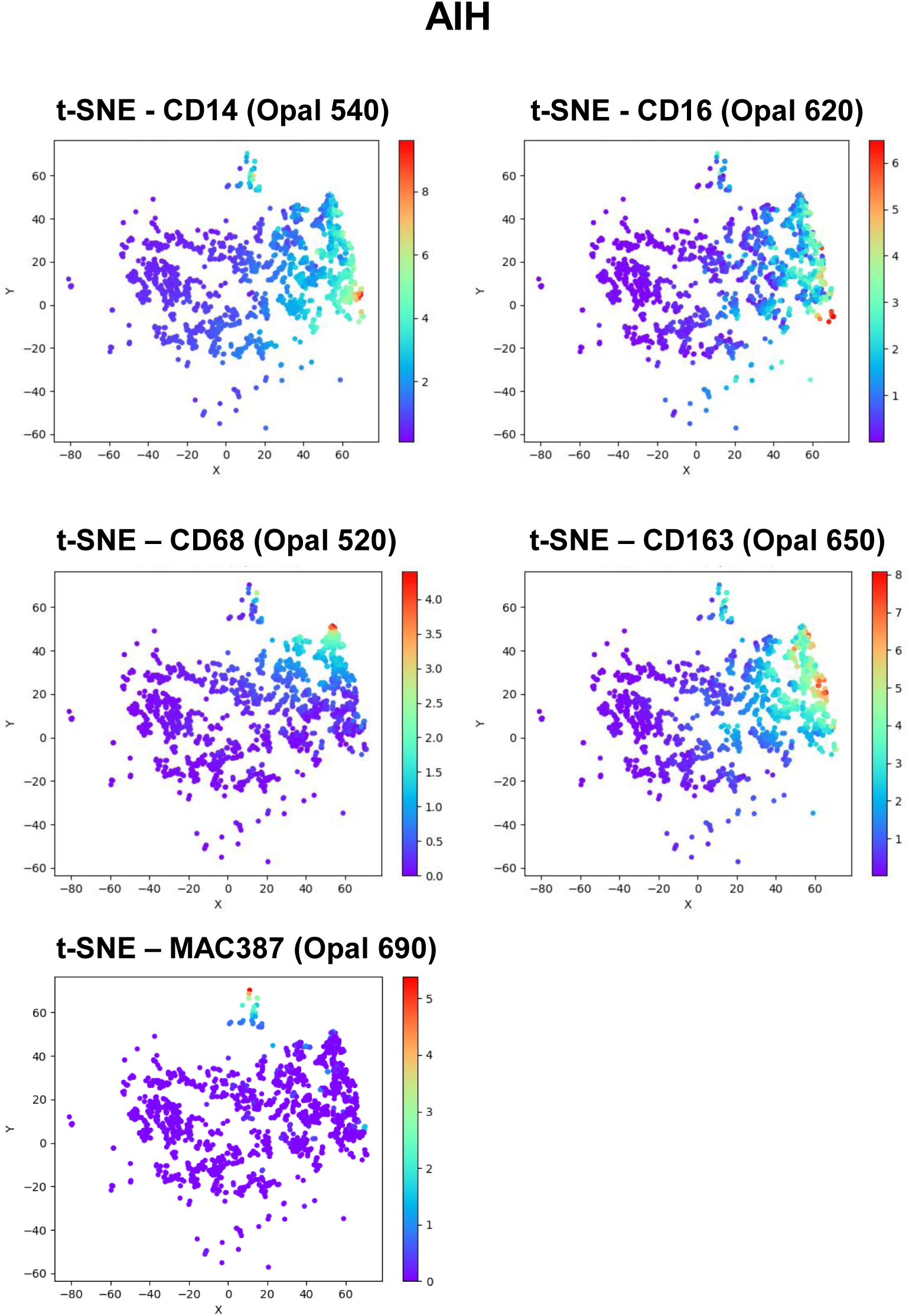
Unsupervised analysis shows increased infiltration of multiple macrophage subpopulations/phenotypes in FFPE liver biopsies obtained from patients with different liver diseases. The t-SNE plots that use dimensional reduction to facilitate visualization of macrophage marker expression, highlight the unique patterns of individual macrophage markers that are present in the livers of patient with chronic liver diseases (**HCV-Suppl Fig 4; NASH-Suppl Fig 5; AIH-Suppl Fig 6**) versus control (Suppl Fig 3). Cells with similar properties appear close together in the two-dimensional map and red (or “hot”) markers show cells with relatively more expression of that specific marker when compared to blue (or “cold”) markers, which indicate absent or minimal expression. Each of the different chronic liver diseases showed more expansion and diversity in cellular phenotypes when compared to the tight clusters observed in controls (also see Fig 6).

## References

1. Crispe IN. Immune tolerance in liver disease. Hepatology 2014;60:2109–2117.

2. Li N, Hua J. Immune cells in liver regeneration. Oncotarget 2017;8:3628–3639.

3. Liaskou E, Zimmermann HW, Li KK, Oo YH, Suresh S, Stamataki Z, Qureshi O, et al. Monocyte subsets in human liver disease show distinct phenotypic and functional characteristics. Hepatology 2013;57:385–398.

4. Demetris AJ, Bellamy CO, Gandhi CR, Prost S, Nakanuma Y, Stolz DB. Functional Immune Anatomy of the Liver-As an Allograft. Am J Transplant 2016;16:1653–1680.

5. McGuinness PH, Painter D, Davies S, McCaughan GW. Increases in intrahepatic CD68 positive cells, MAC387 positive cells, and proinflammatory cytokines (particularly interleukin 18) in chronic hepatitis C infection. Gut 2000;46:260–269.

6. Heymann F, Tacke F. Immunology in the liver--from homeostasis to disease. Nat Rev Gastroenterol Hepatol 2016;13:88–110.

7. MacParland SA, Liu JC, Ma XZ, Innes BT, Bartczak AM, Gage BK, Manuel J, et al. Single cell RNA sequencing of human liver reveals distinct intrahepatic macrophage populations. Nat Commun 2018;9:4383.

8. Soulas C, Conerly C, Kim WK, Burdo TH, Alvarez X, Lackner AA, Williams KC. Recently infiltrating MAC387(+) monocytes/macrophages a third macrophage population involved in SIV and HIV encephalitic lesion formation. Am J Pathol 2011;178:2121–2135.

9. Zimmermann HW, Seidler S, Nattermann J, Gassler N, Hellerbrand C, Zernecke A, Tischendorf JJ, et al. Functional contribution of elevated circulating and hepaticnon-classical CD14CD16 monocytes to inflammation and human liver fibrosis. PLoS One 2010;5:e11049.

10. Tu Z, Pierce RH, Kurtis J, Kuroki Y, Crispe IN, Orloff MS. Hepatitis C virus core protein subverts the antiviral activities of human Kupffer cells. Gastroenterology 2010;138:305–314.

11. Bility MT, Nio K, Li F, McGivern DR, Lemon SM, Feeney ER, Chung RT, et al. Chronic hepatitis C infection-induced liver fibrogenesis is associated with M2 macrophage activation. Sci Rep 2016;6:39520.

12. Zhang Q, Wang Y, Zhai N, Song H, Li H, Yang Y, Li T, et al. HCV core protein inhibits polarization and activity of both M1 and M2 macrophages through the TLR2 signaling pathway. Sci Rep 2016;6:36160.

13. Saha B, Kodys K, Szabo G. Hepatitis C virus-induced monocyte differentiation into polarized M2 macrophages promotes stellate cell activation via TGF-beta. Cell Mol Gastroenterol Hepatol 2016;2:302–316.e308.

14. Guillot A, Tacke F. Liver Macrophages: Old dogmas and new insights. Hepatol Commun 2019;3:730–743.

15. Weston CJ, Zimmermann HW, Adams DH. The role of myeloid-derived cells in the progression of liver disease. Front Immunol 2019;10:893.

16. Li P, He K, Li J, Liu Z, Gong J. The role of Kupffer cells in hepatic diseases. Mol Immunol 2017;85:222–229.

17. Guo LP, Zhou L, Li HX, Zhang J, Wang BM. The study of liver macrophages polarization in patients with autoimmune hepatitis. Zhonghua Nei Ke Za Zhi 2017;56:763–765.

18. Lin R, Zhang J, Zhou L, Wang B. Altered function of monocytes/macrophages in patients with autoimmune hepatitis. Mol Med Rep 2016;13:3874–3880.

19. Malorny U, Neumann C, Sorg C. Influence of various detachment procedures on the functional state of cultured murine macrophages. Immunobiology 1981;159:327–336.

20. Chen S, So EC, Strome SE, Zhang X. Impact of detachment methods on M2 macrophage phenotype and function. J Immunol Methods 2015;426:56–61.

21. Gorris MAJ, Halilovic A, Rabold K, van Duffelen A, Wickramasinghe IN, Verweij D, Wortel IMN, et al. Eight-color multiplex immunohistochemistry for simultaneous detection of multiple immune checkpoint molecules within the tumor microenvironment. J Immunol 2018;200:347–354.

22. Wermuth PJ, Jimenez SA. The significance of macrophage polarization subtypes foranimal models of tissue fibrosis and human fibrotic diseases. Clin Transl Med 2015;4:2.

23. Mezheyeuski A, Bergsland CH, Backman M, Djureinovic D, Sjöblom T, Bruun J, Micke P. Multispectral imaging for quantitative and compartment-specific immune infiltrates reveals distinct immune profiles that classify lung cancer patients. J Pathol 2017.

24. Stack EC, Foukas PG, Lee PP. Multiplexed tissue biomarkerimaging. J Immunother Cancer 2016;4:9.

25. Huang W, Hennrick K, Drew S. A colorful future of quantitative pathology: validation of Vectra technology using chromogenic multiplexed immunohistochemistry and prostate tissue microarrays. Hum Pathol 2013;44:29–38.

26. Faget L, Hnasko TS. Tyramide signal amplification for immunofluorescent enhancement. Methods Mol Biol 2015;1318:161–172.

27. Ishak K, Baptista A, Bianchi L, Callea F, De Groote J, Gudat F, Denk H, et al. Histological grading and staging of chronic hepatitis. J Hepatol 1995;22:696–699.

28. Huang Y, de Boer WB, Adams LA, MacQuillan G, Bulsara MK, Jeffrey GP. Image analysis of liver biopsy samples measures fibrosis and predicts clinical outcome. J Hepatol 2014;61:22–27.

29. Strauss O, Phillips A, Ruggiero K, Bartlett A, Dunbar PR. Immunofluorescence identifies distinct subsets of endothelial cells in the human liver. Sci Rep 2017;7:44356.

30. Uhrig A, Banafsche R, Kremer M, Hegenbarth S, Hamann A, Neurath M, Gerken G, et al. Development and functional consequences of LPS tolerance in sinusoidal endothelial cells of the liver. J Leukoc Biol 2005;77:626–633.

31. van der Maaten L, Hinton G. Visualizing Data using t-SNE. Journal of Machine Learning Research 2008;9:2579–2605.

32. Gadd VL, Melino M, Roy S, Horsfall L, O’Rourke P, Williams MR, Irvine KM, et al. Portal, but not lobular, macrophages express matrix metalloproteinase-9: association with the ductular reaction and fibrosis in chronic hepatitis C. Liver Int 2013;33:569–579.

33. Ju C, Tacke F. Hepatic macrophages in homeostasis and liver diseases: from pathogenesis to novel therapeutic strategies. Cell Mol Immunol 2016;13:316–327.

34. Krenkel O, Tacke F. Liver macrophages in tissue homeostasis and disease. Nat Rev Immunol 2017;17:306–321.

35. Huang R, Wu H, Liu Y, Yang C, Pan Z, Xia J, Xiong Y, et al. Increase of infiltrating monocytes in the livers of patients with chronic liver diseases. Discov Med 2016;21:25–33.

36. Racanelli V, Rehermann B. The liver as an immunological organ. Hepatology 2006;43:S54–62.

37. Sandler NG, Koh C, Roque A, Eccleston JL, Siegel RB, Demino M, Kleiner DE, et al. Host response to translocated microbial products predicts outcomes of patients with HBV or HCV infection. Gastroenterology 2011;141:1220–1230, 1230.e1221-1223.

38. Tippett E, Cheng WJ, Westhorpe C, Cameron PU, Brew BJ, Lewin SR, Jaworowski A, et al. Differential expression of CD163 on monocyte subsets in healthy and HIV-1 infected individuals. PLoS One 2011;6:e19968.

39. Akil A, Endsley M, Shanmugam S, Saldarriaga O, Somasunderam A, Spratt H, Stevenson HL, et al. Fibrogenic gene expression in hepatic stellate cells induced by HCV and HIV replication in a three cell co-culture modelsystem. Sci Rep 2019;9:568.

40. Friedman SL, Ratziu V, Harrison SA, Abdelmalek MF, Aithal GP, Caballeria J, Francque S, et al. A randomized, placebo-controlled trial of cenicriviroc for treatment of nonalcoholic steatohepatitis with fibrosis. Hepatology 2018;67:1754–1767.

41. Reid DT, Reyes JL, McDonald BA, Vo T, Reimer RA, Eksteen B. Kupffer cells undergo fundamental changes during the development of experimental nash and are critical in initiating liver damage and inflammation. PLoS One 2016;11:e0159524.

42. Tacke F. Targeting hepatic macrophages to treat liver diseases. J Hepatol 2017;66:1300–1312.

43. Krenkel O, Puengel T, Govaere O, Abdallah AT, Mossanen JC, Kohlhepp M, Liepelt A, et al. Therapeutic inhibition of inflammatory monocyte recruitment reduces steatohepatitis and liver fibrosis. Hepatology 2018;67:1270–1283.

44. Stack EC, Wang C, Roman KA, Hoyt CC. Multiplexed immunohistochemistry, imaging, and quantitation: a review, with an assessment of Tyramide signal amplification, multispectral imaging and multiplex analysis. Methods 2014;70:46–58.

45. Zaretsky JM, Garcia-Diaz A, Shin DS, Escuin-Ordinas H, Hugo W, Hu-Lieskovan S, Torrejon DY, et al. Mutations associated with acquired resistance to PD-1 blockade in melanoma. N Engl J Med 2016;375:819–829.

46. Liu XD, Hoang A, Zhou L, Kalra S, Yetil A, Sun M, Ding Z, et al. Resistance to antiangiogenic therapy is associated with an immunosuppressive tumor microenvironment in metastatic renal cell carcinoma. Cancer Immunol Res 2015;3:1017–1029.

47. Mlecnik B, Bindea G, Kirilovsky A, Angell HK, Obenauf AC, Tosolini M, Church SE, et al. The tumor microenvironment and Immunoscore are critical determinants of dissemination to distant metastasis. Sci Transl Med 2016;8:327ra326.

48. Knodell RG, Ishak KG, Black WC, Chen TS, Craig R, Kaplowitz N, Kiernan TW, et al. Formulation and application of a numerical scoring system for assessing histological activity in asymptomatic chronic active hepatitis. Hepatology 1981;1:431–435.

